# ERK signaling licenses SKN-1A/NRF1 for proteasome production and proteasomal stress resistance

**DOI:** 10.1101/2021.01.04.425272

**Authors:** Peng Zhang, Hai-Yan Qu, Ziyun Wu, Huimin Na, John Hourihan, Fang Zhang, Feimei Zhu, Meltem Isik, Albertha J.M. Walhout, Yu-Xiong Feng, T. Keith Blackwell

## Abstract

The ubiquitin-proteasome system is vital for cell growth and homeostasis, but for most cancers proteasomal inhibition has not been effective as a therapy. Normal and cancer cells adapt to proteasomal stress through an evolutionarily conserved recovery response, in which the transcription factor NRF1 upregulates proteasome subunit genes. Starting with a *C. elegans* screen to identify regulators of the recovery response, here we show that this response depends upon phosphorylation of NRF1 on a single residue by the growth factor-activated kinase ERK1/2. Inhibition of this phosphorylation impairs NRF1 nuclear localization and proteasome gene activation, sensitizes *C. elegans* and cancer cells to proteasomal stress, and synergizes with proteasome inhibition to retard human melanoma growth *in vivo* in a mouse model. The evolutionarily conserved ERK1/2-NRF1 axis couples proteasome production to growth signaling, and represents a promising new strategy for expanding the range and efficacy of proteasomal inhibition therapy in cancer.

## Introduction

The ubiquitin-proteasome system (UPS) clears damaged or misfolded proteins from the cell, and mediates regulatory mechanisms that involve protein turnover. The proteasome is a highly conserved multiprotein complex that degrades ubiquitin-tagged substrates (Collins and Goldberg, 2017; Tomko and Hochstrasser, 2013). It consists of a barrel-shaped 20S subunit that has multiple proteolytic activities, and is capped on each end by a 19S regulatory subunit that delivers substrates for degradation (Finley, 2009). The proteasome is essential for cellular homeostasis and growth, and organismal development, and its dysfunction has been linked to pathologies that include neurodegenerative diseases and possibly systemic autoimmunity (Ayyadevara et al., 2015; Dantuma and Bott, 2014; Zheng et al., 2016). On the other hand, proteasome inhibition is an important strategy in cancer therapy, and in particular has revolutionized treatment of multiple myeloma, a plasma cell tumor that typically produces high levels of antibody proteins and is under secretory stress (Deshaies, 2014; Guerrero-Garcia et al., 2018; Manasanch and Orlowski, 2017). Unfortunately, however, proteasome inhibitory therapy is ineffective in almost all other cancers, and in multiple myeloma resistance ultimately appears through mechanisms that are poorly understood. Strategies for broadening or prolonging the effectiveness of proteasome-based cancer therapy could represent a major advance.

One way to augment the effectiveness of proteasome inhibition therapy could be to limit proteasome production by the tumor cell. When the proteasome is inhibited in multiple myeloma or other cells, the expression of proteasome subunit genes is increased (Fleming et al., 2002; Meiners et al., 2003; Mitsiades et al., 2002; Wójcik and DeMartino, 2002). This increase is mediated by an evolutionarily conserved transcriptional “recovery” response that is orchestrated by the transcription factor NRF1 (NF-E2-related factor 1, or NFE2L1), and boosts proteasome levels by directly upregulating expression of essentially all proteasome subunit genes (Lehrbach and Ruvkun, 2016; Li et al., 2011; Radhakrishnan et al., 2010; Steffen et al., 2010). In mice NRF1 is essential for development, but tissue-specific knockout studies have demonstrated that it is needed to prevent neurodegeneration, and is important for lipid and cholesterol metabolism in the liver (Kobayashi et al., 2011; Lee et al., 2011; Tsujita et al., 2014; Widenmaier et al., 2017; Xu et al., 2005). Importantly, cultured mammalian cells become sensitized to proteasome inhibition when NRF1 is depleted (Acosta-Alvear et al., 2015; Radhakrishnan et al., 2010; Sha and Goldberg, 2014), suggesting that pharmacological targeting of the proteasome recovery response might be a promising strategy for enhancing the effectiveness of proteasomal inhibition therapy.

A mechanistic understanding of NRF1 regulation could potentially be valuable for intervention in cancer or other disease settings. NRF1 is controlled through a complex process in which it is translated into the endoplasmic reticulum (ER) lumen, where it becomes N-glycosylated and is inserted into the ER membrane at its N-terminus (Deshaies, 2014; Zhang et al., 2006) (Figure 1A). Within the ER NRF1 is perceived to be an unfolded protein, and is extruded into the cytoplasm by the ER-associated degradation (ERAD) mechanism, so that NRF1 becomes ubiquitinated and ultimately is degraded by the proteasome (Radhakrishnan et al., 2014; Sha and Goldberg, 2014). NRF1 becomes stabilized when proteasome activity is insufficient or inhibited, and thereby senses proteasomal stress directly (Radhakrishnan et al., 2014; Radhakrishnan et al., 2010; Sha and Goldberg, 2014; Steffen et al., 2010). In addition, NRF1 transcriptional activity and the proteasome recovery response depend upon NRF1 being de-N-glycosylated (deglycosylated) by the cytoplasmic N-glycanase NGLY1, cleaved near its N-terminus by the serine protease DDI2/DDI-1, and stabilized by attachment of O-linked N-acetylglucosamine (O-GlcNAc) (Han et al., 2017; Koizumi et al., 2016; Sekine et al., 2018; Tomlin et al., 2017) (Figure 1A).

**Figure 1.**
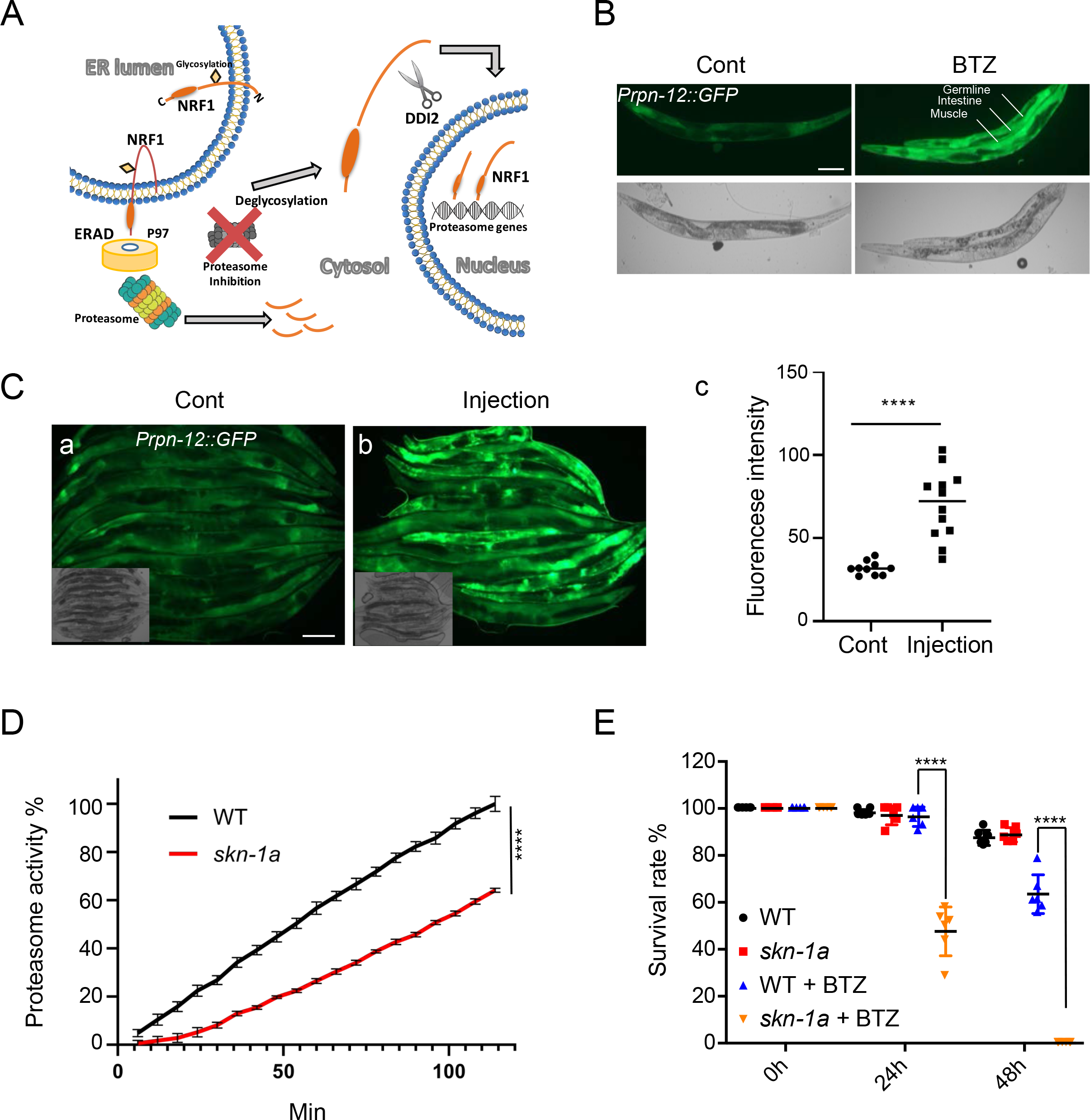
SKN-1A mediates a conserved proteasome recovery response to BTZ. **A**. Features of the NRF1-mediated proteasome recovery response that are critical background for this study. **B**. BTZ activates the proteasome subunit transcriptional reporter *Prpn-12::GFP* in the intestine, muscle and germline. Scale bar, 100 µm. **C**. Induction of the proteasome recovery response by injection of 200 nM BTZ into *C. elegans*. Quantification and statistics quantification are shown in the right panel c, Scale bar, 100 µm, ****p<0.0001. **D**. In *skn-1a* mutant worms, chymotrypsin-like proteasome activity is lower than in the wild type, ****p<0.0001. **E**. The *skn-1a* reduces resistance to 500 µM BTZ-induced proteasomal stress, ****p<0.0001.

This complex mode of NRF1 regulation appears to be largely conserved in the nematode *C. elegans*, in which a single orthologous gene (*skn-1*) (Bowerman et al., 1992) encodes alternatively spliced isoforms that correspond to NRF1 (SKN-1A) and the related transcription factor NRF2 (SKN-1C) (Blackwell et al., 2015) (Figure 1A and S1A). NRF2/SKN-1C, which has been studied far more extensively than NRF1, mediates the major response to oxidative stress and is regulated within the cytoplasm, not the ER (Blackwell et al., 2015; Suzuki and Yamamoto, 2015). In *C. elegans skn-1* is required for maintenance of steady-state proteasome activity, indicating that this is an important homeostasis mechanism, and for essentially all proteasome genes to be upregulated in response to stress (Li et al., 2011; Oliveira et al., 2009). These functions are mediated by the NRF1-like SKN-1A isoform (Lehrbach and Ruvkun, 2016), which differs from SKN-1C only in the presence of 90 additional residues at its N-terminus that contain a transmembrane segment (Figure S1A) (Lehrbach and Ruvkun, 2016). Mutational inactivation of SKN-1A/NRF1 sensitizes *C. elegans* to developmental arrest induced by the proteasome inhibitor Bortezomib (BTZ) (Lehrbach and Ruvkun, 2016; Lehrbach and Ruvkun, 2019), a first-line therapy for multiple myeloma (Guerrero-Garcia et al., 2018), and SKN-1A is required for proteasomal stress to activate a reporter gene in which the promoter for a proteasomal subunit gene (*rpt-3*) is fused to green fluorescent protein (GFP) (Lehrbach and Ruvkun, 2016). Like NRF1, SKN-1A localizes to the ER (Glover-Cutter et al., 2013), and for its transcriptional activity requires glycosylation in the ER, ERAD functions, deglycosylation by NGLY1, and cleavage by DDI2/DDI-1 (Lehrbach et al., 2019; Lehrbach and Ruvkun, 2016; Lehrbach and Ruvkun, 2019). These parallels suggest that fundamental features of the proteasome recovery response have been maintained during approximately one billion years of evolution.

Here we used genome-scale RNA interference (RNAi) screening in *C. elegans* to uncover mechanisms that are required for the SKN-1A/NRF1-mediated proteasome recovery response *in vivo*. We found that full-scale activity of this response and resistance to proteasomal stress depends upon MPK-1, the *C. elegans* ortholog of ERK1/2, which is the canonical kinase target of RAS-RAF-MEK signaling in growth factor responses (Roux and Blenis, 2004). In human cells, NRF1 interacts with and is phosphorylated by ERK1/2.

Phosphorylation by ERK1/2 is required for NRF1 nuclear localization, and the proteasome recovery response. Pharmacological inhibition of ERK1/2 sensitizes human cancer cells to proteasomal inhibition, and simultaneous ERK1/2 and proteasome inhibition synergistically inhibits the growth of transplanted human melanoma cells *in vivo* in mice. Our findings suggest that growth signaling through ERK1/2 functions as an evolutionarily conserved licensing mechanism for NRF1 regulation of proteasome expression, and identify the ERK1/2-NRF1 pathway as a promising novel target for pharmacologically enhancing the therapeutic efficacy of proteasome inhibition in cancer therapy.

## Results

### SKN-1A mediates a conserved proteasome recovery response to BTZ

Prior to studying the proteasome recovery response through *C. elegans* screening, we further investigated the extent to which this response resembles that of mammals. A reporter in which the *rpn-12* proteasome subunit gene promoter regulates GFP expression (*Prpn-12::GFP*) was robustly activated in the intestine, muscle and germline by BTZ treatment but not by various other stresses (Figure 1B, S1B). BTZ treatment also resulted in accumulation of the SKN-1A protein in nuclei in adult animals (Figure S1C). While these data are consistent with evolutionary conservation of the proteasome recovery response, in *C. elegans* much higher BTZ concentrations are required to induce toxicity (Lehrbach and Ruvkun, 2016), or activation of SKN-1A/NRF1 and proteasome genes (Figure S1D), than are needed in cultured mammalian cells (1-100 µM compared to 1-200 nM) (Adams et al., 1999; Lehrbach and Ruvkun, 2016; Lehrbach and Ruvkun, 2019). This difference is typical of drugs that do not cross the *C. elegans* cuticle and must be ingested (Page et al., 2014), but it nevertheless raises the possibility that the *C. elegans* proteasome might be relatively insensitive to BTZ, and thus that BTZ toxicity could involve off-target effects. Importantly, however, addition of 100 nM BTZ directly to a *C. elegans* lysate saturated the proteasome chymotrypsin activity (Figure S1E), and injection of 200 nM BTZ into the *C. elegans* gut activated *Prpn-12::GFP* (Figure 1C). This indicates that *C. elegans* is comparable to mammals with respect to the sensitivity of the proteasome, and the proteasome recovery response to BTZ treatment. Previously, SKN-1A nuclear localization and proteasome subunit genes were also activated by RNAi knockdown of other proteasome genes, another mode of proteasome activity inhibition (Lehrbach and Ruvkun, 2016; Li et al., 2011). Furthermore, in adult animals a mutation that specifically eliminates SKN-1A (*mg570*) (Figure S1A) (Lehrbach and Ruvkun, 2016) dramatically reduced proteasome activity (Figure 1D) and compromised resistance to BTZ-induced death (Figure 1E). Together, the data support the idea that *C. elegans* is an appropriate *in vivo* model for investigating the effects of BTZ on the proteasome recovery response.

If the *C. elegans* proteasome recovery response is analogous to that of mammals (Radhakrishnan et al., 2010; Sha and Goldberg, 2014; Steffen et al., 2010), BTZ treatment should activate essentially all proteasome subunit genes through SKN-1A. To investigate whether SKN-1A is generally required for proteasome subunit gene expression in *C. elegans*, we performed RNA-sequencing (RNAseq) transcriptome analysis of wild-type (WT) and *skn-1a* mutant animals under control (M9 buffer) and BTZ conditions (Figure 2A–2D and S2A, B; Table S2 and S3). Most genes that were activated or downregulated by SKN-1A were induced or downregulated, respectively, by BTZ (Figure 2B and 2C), and BTZ treatment induced SKN-1A-dependent activation of all detectable proteasome subunit genes (Figure 2D). These expected results emphasize the critical importance of SKN-1A for proteasome recovery under proteasomal stress conditions, and support the validity of our transcriptome data overall. In addition to its key function in proteasome maintenance (Radhakrishnan et al., 2014; Radhakrishnan et al., 2010; Sha and Goldberg, 2014; Steffen et al., 2010) NRF1 regulates other cellular functions including stress responses (Ohtsuji et al., 2008), embryonic development (Chan et al., 1998; Chen et al., 2003), inflammatory responses (Xu et al., 2005), metabolism (Zhang et al., 2014), and cholesterol homeostasis (Widenmaier et al., 2017). A variety of functions were also represented among genes that are dependent upon SKN-1A (Figure 2A and Table S3). The SKN-1A-upregulated genes we identified overlapped only to a limited extent with SKN-1-upregulated genes identified previously (Ewald et al., 2015; Steinbaugh et al., 2015) (Figure S2B), possibly because those studies involved loss of all SKN-1 isoforms and different conditions. Together, our results indicate that the major features of the NRF1/SKN-1A-mediated proteasome recovery response are evolutionarily conserved.

**Figure 2.**
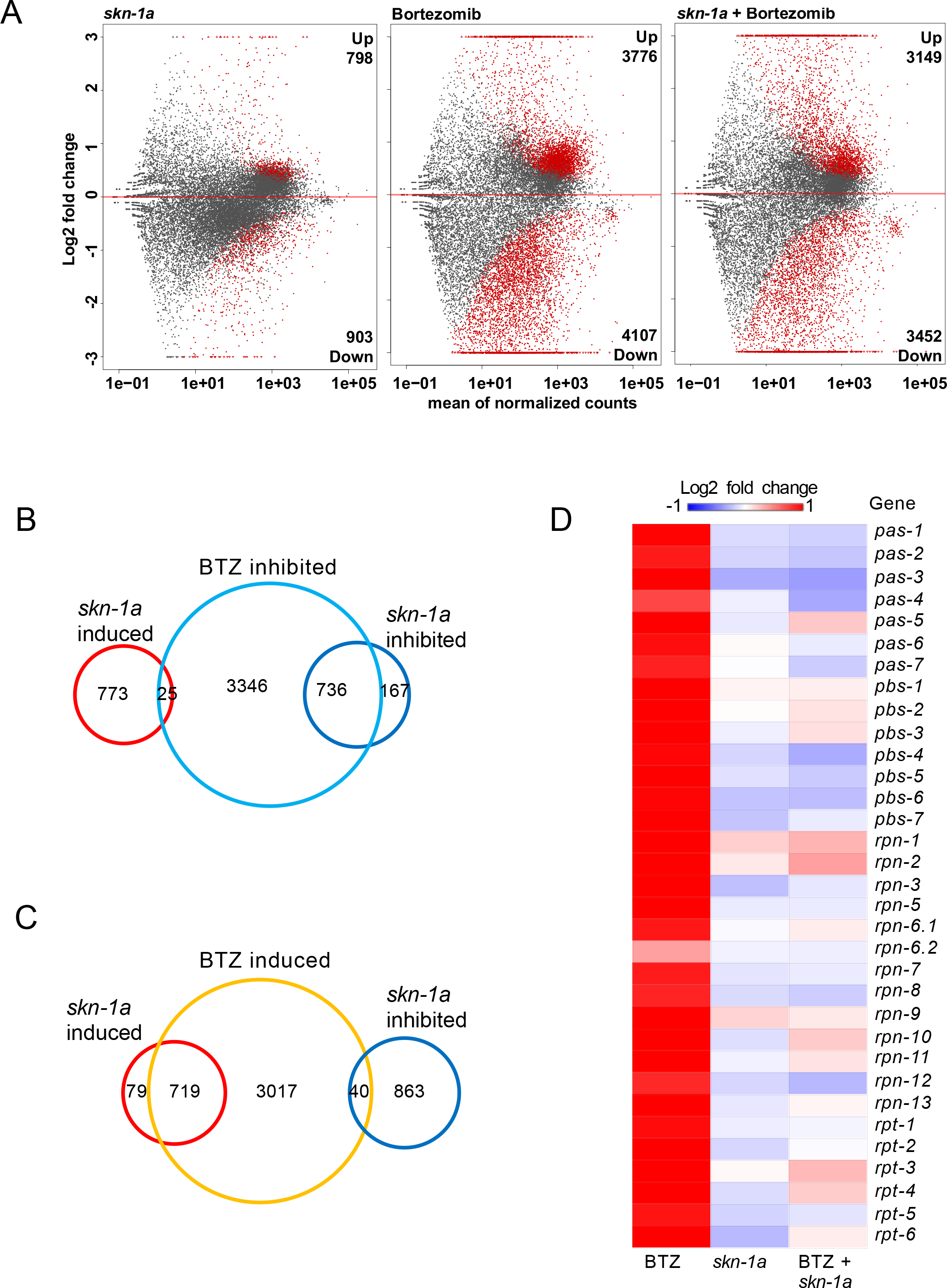
Regulation of proteasome subunit and multiple other gene categories by SKN-1A and BTZ. **A**. MA plot showing analysis of differentially expressed (DE) genes. Red points represent DE genes that differ in expression with a FDR (false discovery rate) of < 0.01. **B, C**. Comparison of genes that are induced or inhibited by BTZ or *skn-1a*. Note that BTZ and SKN-1A act largely in the same direction **D**. Heatmap of log2 fold changes for 33 proteasome subunit genes that are upregulated by BTZ treatment in WT *C. elegans*.

### Dependence of the *C. elegans* proteasome recovery response on MPK-1/ERK signaling

To identify mechanisms that are required for the proteasome recovery response to occur, we screened a library of 11,560 genes by RNAi in *C. elegans* (Rual et al., 2004) to identify those for which knockdown impaired or blocked activation of *Prpn-12::GFP* by BTZ (Figure 3A). This screen identified 124 candidate genes as being needed for this response (Table S1). In mammalian cells the proteasome recovery response depends upon the p97/VCP ATPase, which is required for NRF1 to be relocalized from the ER lumen to the cytosol through the ERAD mechanism (Figure S4C) (Radhakrishnan et al., 2014). Our *C. elegans* screen identified *cdc-48*.*1* and *cdc-48*.2, the *C. elegans* p97 orthologs along with the p97 complex member *ufd-1*, and *skn-1* itself (Table S1). These findings support the validity of our screen, and further indicate the evolutionary conservation of the recovery response.

**Figure 3.**
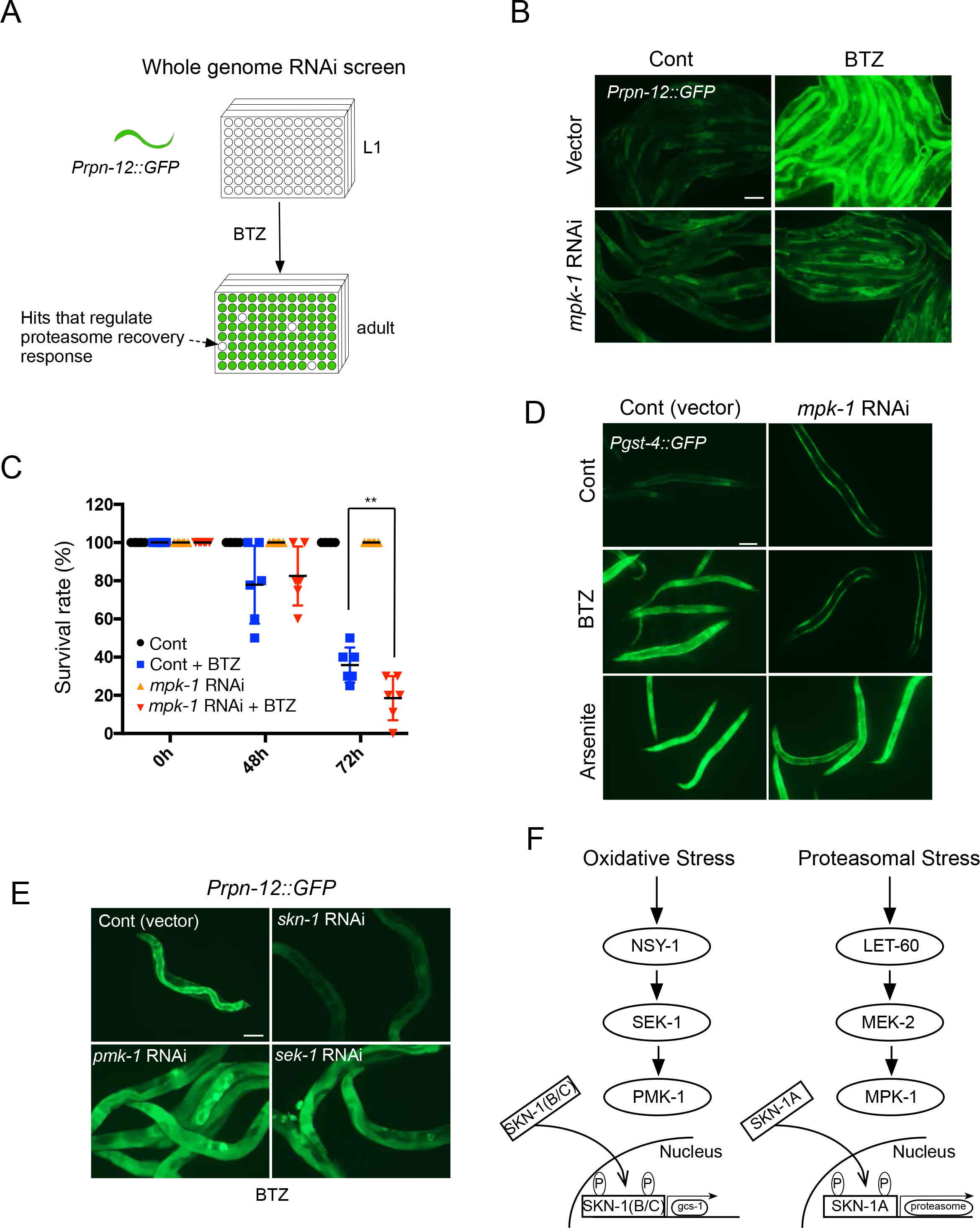
Dependence of the *C. elegans* proteasome recovery response on MPK-1/ERK signaling. **A**. Genome-scale RNAi screening for genes that are required in the proteasome recovery response. In *C. elegans*, RNAi can be imposed by feeding of bacteria that express individual dsRNA clones of interest (Kamath and Ahringer, 2003). **B**. *mpk-1* RNAi blocks the *Prpn-12::GFP* proteasome recovery response induced by BTZ, Scale bar, 100 µm. **C**. Under proteasome inhibition conditions, *C. elegans* dies comparably rapidly after knockdown of the *mpk-1* or s*kn-1* genes, **p<0.01. **D**. In *mpk-1(RNAi)* worms, *Pgst-4::GFP* is activated by Arsenite (oxidative stress) but not BTZ, Scale bar, 100 µm. **E**. In *pmk-1* and *sek-1* mutants, the *Prpn-12::GFP* recovery response is not affected, Scale bar, 100 µm. **F**. A schematic diagram of the PMK-1/SKN-1C and MPK-1/SKN-1A stress response pathways. *gcs-1* is a canonical p38- and oxidative stress-induced target of SKN-1 (Blackwell et al., 2015; Inoue et al., 2005).

We were intrigued that our screen detected the *C. elegans* ERK1/2 ortholog MPK-1 (Table S1, Figure 3B), because the *C. elegans* oxidative stress response that is mediated by SKN-1C/NRF2 is activated by signaling through the MAPK p38, which is related to MPK-1/ERK1/2 (Figure 3C) (Hourihan et al., 2016; Inoue et al., 2005). If p38 phosphorylation of SKN-1 is prevented by mutation or genetic kinase ablation, SKN-1C nuclear accumulation and this oxidative stress response fail to occur. Interestingly, MPK-1 can phosphorylate a SKN-1 protein *in vitro* on the same two amino acids at which SKN-1C is phosphorylated by the *C. elegans* p38 kinase (PMK-1, 2, 3) (Okuyama et al., 2010), but the possible function of SKN-1 phosphorylation by MPK-1 has remained unknown. In light of these earlier findings, detection of *mpk-1* in our screen suggested an interesting model: that because the oxidative and proteasome recovery responses are mediated by different SKN-1 isoforms, these responses might depend specifically upon signaling to SKN-1 through the p38 or MPK-1/ERK pathways, respectively (Figure 3F).

Supporting the importance of MPK-1 in the proteasome recovery response, RNAi against its upstream kinase MEK-2 (Figure S3A) also impeded induction of *Prpn-12::GFP* by BTZ, and knockdown of *mpk-1* comparably sensitized adult *C. elegans* to toxicity from BTZ (Figure 3C). The well-characterized SKN-1 target *gst-4*, which is upregulated in the p38/SKN-1C-mediated oxidative stress response (Glover-Cutter et al., 2013; Hourihan et al., 2016; Inoue et al., 2005), was also activated by BTZ (Figure 3D and S3B). Remarkably, *mpk-1* RNAi inhibited *gst-4* upregulation by BTZ, but not by oxidative stress (Figure 3D, S3B and S3C).

Mirroring this finding, genetic inhibition of p38 signaling blocks *gst-4* activation by oxidative stress (Glover-Cutter et al., 2013), but not *Prpn-12::GFP* activation by BTZ (Figure 3E). We conclude that the SKN-1C-mediated oxidative stress response and the SKN-1A-mediated proteasome recovery response specifically depend upon SKN-1 phosphorylation by the p38 and MPK-1/ERK kinases, respectively (Figure 3F).

### Human NRF1 nuclear localization depends upon phosphorylation by ERK1/2

In mammals, the NRF1-related transcription factor NRF2 is regulated by at least two evolutionarily-conserved phosphorylation events. Both NRF2 and *C. elegans* SKN-1C are activated in response to oxidative stress-induced phosphorylation by p38 (Blackwell et al., 2015; Hourihan et al., 2016; Inoue et al., 2005; O’Connell and Hayes, 2015; Suzuki and Yamamoto, 2015), and inhibited through phosphorylation by glycogen synthase kinase-3 (GSK-3) (Blackwell et al., 2015; Cuadrado, 2015). NRF1 has been reported to be phosphorylated on serine residues, but the regulatory significance of these events has remained largely unclear (Chepelev et al., 2011; Tsuchiya et al., 2013). Given the evolutionary conservation of other mechanisms that regulate NRF1 in the proteasome recovery response, our *C. elegans* findings suggested that mammalian NRF1 might be phosphorylated by ERK1/2. To test this idea we investigated whether ERK1/2 and NRF1 interact physically. As described previously (Radhakrishnan et al., 2014; Radhakrishnan et al., 2010), FLAG-tagged NRF1 that was expressed in HEK293 cells was stabilized by treatment with BTZ (Figure S4D). This NRF1 protein interacted with endogenous ERK1/2, mainly with the 42 kDa form of ERK2, as detected by co-immunoprecipitation with an antibody against ERK1/2 (Figure 4A). Higher levels of NRF1-bound ERK1/2 were isolated from BTZ-treated cells (Figure 4A), which could derive from the increased levels of NRF1 expression (Figure S4D) or ERK1/2 phosphorylation (Figure S5E). Treatment with a pharmacological ERK1/2 inhibitor (SCH772894 or SCH772894-HCL, Figure S5E) decreased the levels of ERK1/2 that interacted with NRF1 either with or without BTZ treatment, indicating that this interaction is dependent upon the ERK1/2 phosphorylation activity.

**Figure 4.**
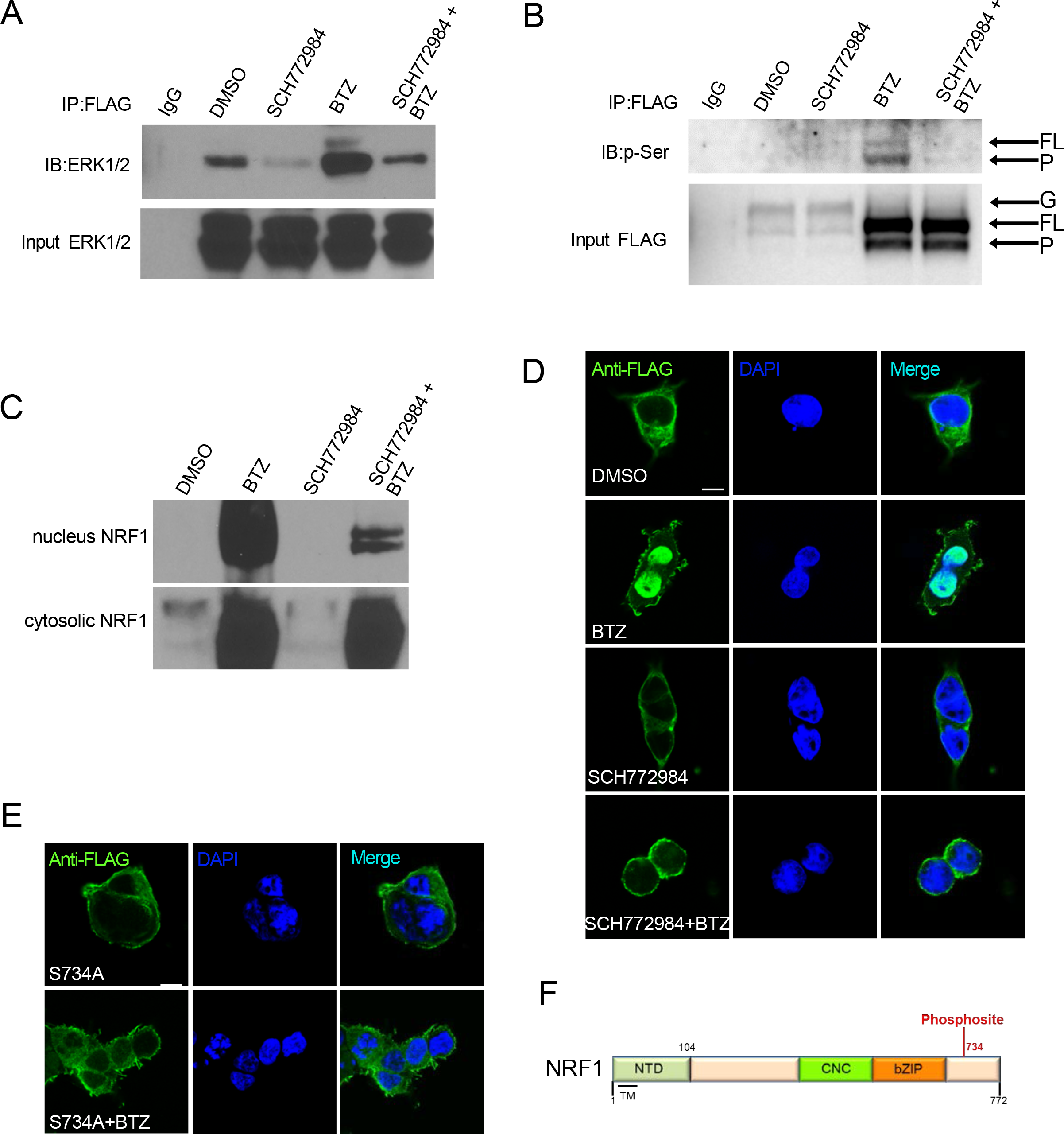
Human NRF1 nuclear localization depends upon phosphorylation by ERK1/2. **A**. Interaction between NRF1 and ERK1/2. Co-immunoprecipitation from HEK293 cells revealed a physical interaction between ERK1/2 and NRF1^3XFLAG^ that is enhanced by BTZ and inhibited by SCH772984. **B**. Dependence of NRF1 serine phosphorylation on ERK1/2. Immunoprecipitation from HEK293 cells showing that phosphorylation of NRF1^3XFLAG^ was blocked by SCH772984. **C**. BTZ-induced nuclear localization of NRF1 is dependent upon ERK1/2. A western blot shows that treatment decreases representation of NRF1 in the nuclear fraction. **D**. Immunofluorescence staining showing that SCH772984 inhibits BTZ-induced accumulation of NRF1^3XFLAG^ protein in HEK293 cell nuclei, which are labeled with DAPI. Quantification is shown in Figure S4F. Scale bar, 10 µm. **E**. The phosphosite S734A is required for NRF1 nuclear localization. Immunofluorescence staining and analysis was performed as in D. Scale bar, 10 µm. **F**. Diagram indicating phosphosite S734 within NRF1.

In mammalian cells NRF1 is detectable in three major species, the relative abundance of which varies according to conditions: a 130 KD glycosylated form (G), a 120 KD full length (FL) species that has been deglycosylated in the cytoplasm by NGLY1, and a deglycosylated 110 KD species that has been processed (P) by DDI2 (Koizumi et al., 2016; Lehrbach and Ruvkun, 2016; Radhakrishnan et al., 2014; Radhakrishnan et al., 2010; Sha and Goldberg, 2014; Tomlin et al., 2017) (Figure S4D). The levels of NRF1, particularly of the cleaved form, are increased dramatically when the proteasome is inhibited (Figure S4D). These forms can be detected with antibodies against endogenous NRF1, or when a FLAG-tagged NRF1 protein is expressed from a cDNA (Figure S4A, D) (Radhakrishnan et al., 2014; Radhakrishnan et al., 2010). FLAG-tagged NRF1 that was immunoprecipited from BTZ-treated HEK293 cells was readily detected by an anti-phosphoserine antibody (Figure 4B). This signal was not detectable in the absence of BTZ treatment, possibly because the levels of NRF1 were far lower. Interestingly, we also did not detect phosphorylation of the glycosylated form of NRF1, and phosphorylation of the FL form was proportionally lower than observed for the processed form (Figure 4B). This suggests that NRF1 becomes phosphorylated primarily after deglycosylation and cleavage.

Importantly, the NRF1 phosphorylation signal was largely eliminated in cells that were treated with an ERK1/2 inhibitor (Figure 4B), suggesting that this phosphorylation is mediated largely by ERK1/2. Mass spectrometry (MS) phosphoproteomic analysis of FLAG-tagged NRF1 did not detect any phosphopeptides in untreated cells, but detected five phosphopeptides in BTZ-treated cells, each of which included a consensus ERK1/2 phosphorylation motif PYSP at S734 (Figure S4B-S4E). This difference parallels the increase in serine phosphorylation of NRF1 that we detected by western blotting (Figure 4B), and presumably derives from the increases in NRF1 levels (Figure S4D) and possibly ERK1/2 activity (Figure S5E) that are induced by BTZ treatment. By contrast, only one phosphopeptide (at S734) was detected in the sample that was treated with an ERK1/2 inhibitor (not shown). This apparent ERK1/2 phosphorylation site is distinct from the positions at which NRF1 has previously been reported to be phosphorylated (Okuyama et al., 2010)). Taken together, these data suggest that NRF1 is phosphorylated by ERK1/2 at S734 (Figure 4F), and that this residue represents its most prominent phosphorylation site.

Pharmacological inhibition of ERK1/2 did not detectably impede NRF1 processing or the increase in NRF1 levels that follows from proteasome inhibition (Figure S4A, D), but interfered with the localization of NRF1 to nuclei (Figure 4C). Analysis of cellular nuclear and cytoplasmic fractions revealed a dramatic increase in NRF1 levels associated with BTZ treatment, and accumulation of the full-length and processed but not glycosylated forms in nuclei (Figure 4C), as reported previously (Radhakrishnan et al., 2014). Inhibition of ERK1/2 did not alter the accumulation of NRF1 forms within the cytoplasm (Figure 4C), consistent with its phosphorylation of NRF1 occurring largely after NRF1 is N-deglycosylated and cleaved (Figure 4B). However, ERK1/2 inhibition dramatically reduced the levels of NRF1 in nuclei (Figure 4C). Supporting this idea, immunofluorescence analysis of FLAG-tagged NRF1 also indicated that ERK1/2 inhibition inhibited the nuclear accumulation of NRF1 that otherwise results from BTZ treatment (Figure 4D and S4F). Similarly, a mutant version of NRF1 in which the ERK1/2 phosphorylation site at S734 had been changed to alanine largely failed to accumulate in nuclei in response to BTZ treatment (Figure 4E and S4F). We conclude that phosphorylation of NRF1 by ERK1/2 important for BTZ-induced localization of NRF1 to the nucleus, but is dispensable for upstream NRF1 processing steps.

### ERK1/2 inhibition sensitizes human tumor cells to proteasomal stress

Considering the importance of both SKN-1A and the ERK1/2 kinase MPK-1 for resistance to proteasomal stress in *C. elegans* (Lehrbach and Ruvkun, 2016) (Figure 1E and 3C), we investigated the extent to which these mechanisms contribute to proteasome function and proteasomal stress resistance in cultured mammalian tumor cells. In accordance with SKN-1A being required for *C. elegans* proteasome activity, in human SH-SY5Y neuroblastoma cells that express shRNA to NRF1 (Sha and Goldberg, 2014) the steady-state Chymotrypsin-like proteasome activity was reduced (Figure S6A). As reported previously (Sha and Goldberg, 2014), knockdown of NRF1 dramatically sensitized SH-SY5Y cells to BTZ treatment, as indicated by increased levels of an apoptosis protein marker, cleaved Caspase3 (Porter, 1999) (Figure 5A). Knockdown of NRF1 but not NRF2 similarly sensitized human HepG2 liver carcinoma cells to BTZ treatment, consistent with evidence that the mammalian and *C. elegans* proteasome recovery responses are mediated by NRF1/SKN-1A but not NRF2/SKN-1C (Lehrbach et al., 2019; Radhakrishnan et al., 2014; Radhakrishnan et al., 2010) (Figure 5B). Together, these findings indicate that interference with NRF1 function might be a promising strategy for enhancing the effectiveness of proteasome inhibition therapy in cancer.

**Figure 5.**
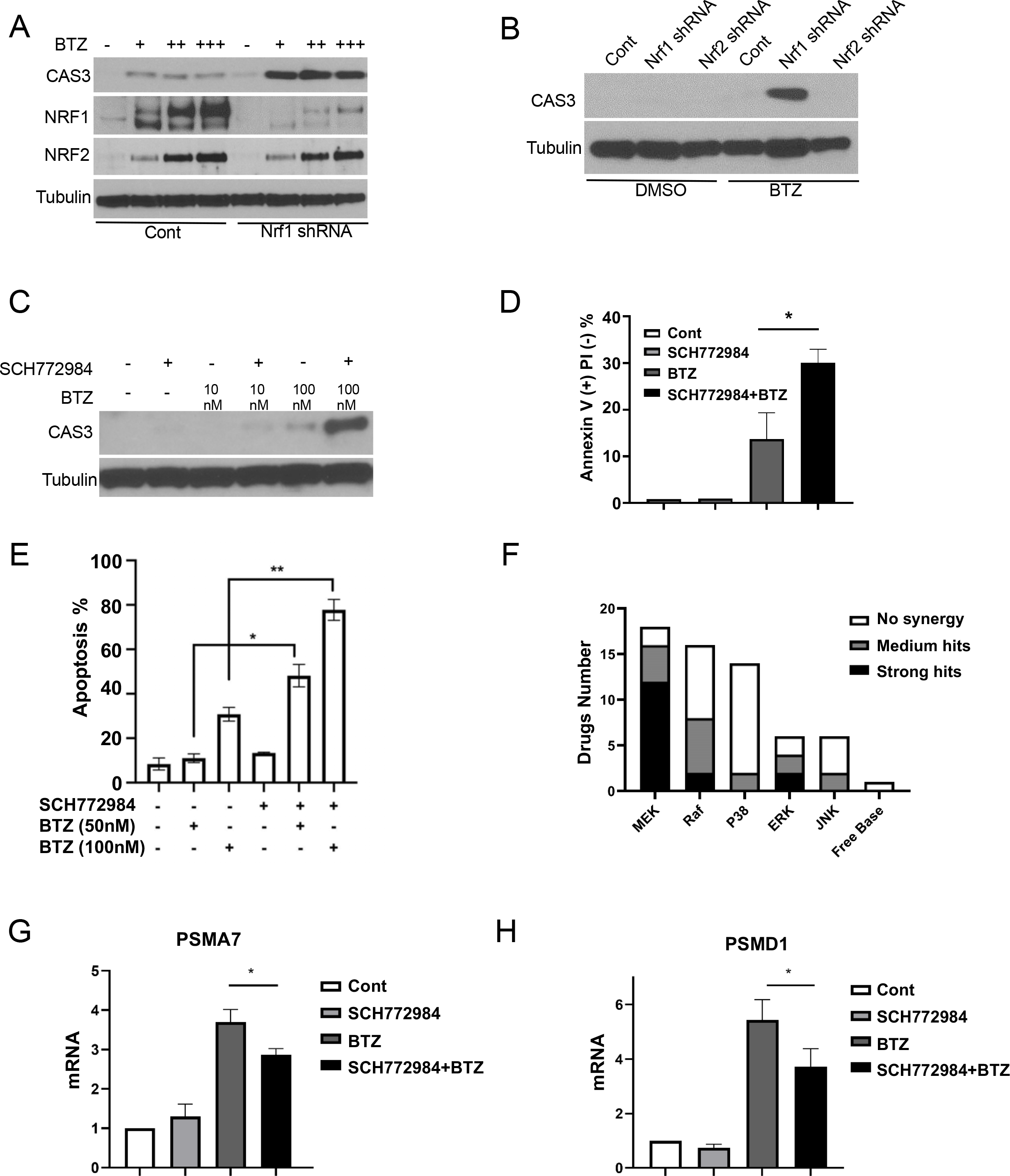
ERK1/2 inhibition sensitizes human tumor cells to proteasomal stress. **A**. NRF1 knockdown sensitizes tumor cells to BTZ-induced cell death. Western blotting shows that NRF1 knockdown increases BTZ-induced cell death in HepG2 cells, indicated by CAS3 cleavage. **B**. NRF1 but not NRF2 knockdown sensitizes HepG2 cells to BTZ-induced apoptosis. **C**. Inhibition of ERK1/2 by SCH772984 sensitizes HepG2 cells to BTZ-induced apoptosis. **D**. Inhibition of ERK1/2 by SCH772984 sensitizes MM.1s multiple myeloma cells to BTZ-induced apoptosis. Propidium iodide and Annexin V staining were monitored by flow cytometry. *p<0.05. **E**. SCH772984 sensitizes A2058 melanoma cells to BTZ-induced apoptosis, detected by flow cytometry, *p<0.05, **p<0.01. **F**. Result of a small inhibitor screen to identify drugs that increase HepG2 cell death synergistically with BTZ, detected by CAS3 western blot. 16 of 18 MEK inhibitors and 4 of 6 ERK inhibitors showed a positive synergistic effect. **G, H**. BTZ-induced expression of *psma7* (**G**) and pcmd1 (**H**) proteasome subunit genes is decreased by SCH772894 treatment, assayed by RT-PCR in HepG2 cells, *p<0.05.

While transcription factors in general represent challenging targets for pharmacological intervention, our observations suggest that targeting of ERK1/2 might be a promising strategy for interfering with the proteasome recovery response therapeutically. Multiple myeloma cells are extremely sensitive to proteasome inhibition (Adams, 2004; Gandolfi et al., 2017), but despite being under secretory stress these cells attempt to increase proteasome levels when challenged with a proteasome inhibitor (Mitsiades et al., 2002). This suggests that the proteasome recovery response might promote multiple myeloma cell growth and survival. Supporting this idea, ERK1/2 inhibition synergized with BTZ treatment to reduce proliferation or viability of the multiple myeloma cell line MM.1s (Figure S5F). Additionally, two apoptosis assays indicated that ERK1/2 inhibition dramatically increased the level of BTZ-induced killing of MM.1s (Figure 5D and S5C, D). Thus, inhibition of ERK1/2 and the proteasome recovery response enhanced the already highly effective killing of multiple myeloma cells by BTZ.

Solid tumors represent a more challenging target for proteasome intervention therapy, with 10-50 fold higher concentrations of BTZ being needed to induce cell death than is typical for multiple myeloma cells (Adams, 2004). Importantly, however, we determined that in HepG2 and human A2058 melanoma cells pharmacological ERK1/2 inhibition did not induce cell death on its own, but dramatically increased the extent to which BTZ treatment triggered apoptosis (Figure 5C-E). To examine the relative importance of ERK1/2 signaling for resistance to BTZ, and to test more than one compound that acts on this pathway, we screened HepG2 cells with a small library of MAP kinase pathway inhibitors to identify compounds and pathways that are associated with BTZ resistance (Figure 5F and Figure S6C-E). Inhibitors of p38 signaling largely failed to increase cell death in the presence of BTZ (Figure 5F), consistent with our evidence that this pathway is dispensable for the SKN-1A-mediated proteasome recovery response in *C. elegans* (Figure 3F). By contrast, and consistent with the importance of ERK1/2 signaling for NRF1 function, most inhibitors of Ras-MEK-ERK pathway components (Raf, MEK, ERK) increased BTZ-associated cell death under conditions where they did not trigger apoptosis as single agents (Figure 5F). Pharmacological inhibition of ERK1/2 did not detectably reduce the steady-state level of proteasome activity (Figure S6B), but did slow the upregulation of proteasomal subunit genes that is characteristic of the proteasome recovery response (Figure 5G, H). The latter result would be expected, given that phosphorylation of NRF1 by ERK1/2 is important for BTZ-induced NRF1 nuclear localization (Figure 4C-E). We conclude that ERK1/2 phosphorylation of NRF1 is critical for the proteasome recovery response, and that by impairing this response ERK1/2 inhibition can sensitize a variety of different human tumor cell types to BTZ toxicity.

### Synergistic inhibition of melanoma growth by proteasome and ERK1/2 inhibition *in vivo*

Encouraged by our findings, we investigated whether ERK1/2 inhibition might sensitize A2058 melanoma cells to BTZ treatment in a preclinical mouse model. Consistent with their moderate response to BTZ *in vitro* (Figure 5E and S5A, B), the growth of A2058 cells was only mildly retarded by BTZ treatment *in vivo* (Figure 6A). Importantly, the tumor-inhibitory effect of BTZ was dramatically increased when tumors were co-treated with the ERK1/2 inhibitor SCH772984. While SCH772984 alone did not alter tumor growth, it synergized with BTZ to suppress the growth of A2058 tumors (Figure 6A, Combination index: 0.59). The tumor growth inhibitory (TGI) effect of BTZ was increased by over 2-fold when SCH772984 was co-administered (Figure 6B). Staining for Ki67, a mitotic marker, in the A2058 tumors revealed that proliferation of cancer cells was inhibited by combination treatment of SCH772984 and BTZ compared to BTZ alone (Figure 6C, D), consistent with the change in tumor weight (Figure 6E, Figure S7A). Importantly, treatment with SCH772984 dramatically inhibited the phosphorylation of ERK1/2, while sparing MEK1/2, suggesting that the synergistic anti-tumor effect of SCH772984 was mediated by inhibition of ERK1/2 but not MEK1/2 (Figure S7B). Mechanistically, we found that the proteasome recovery response that was induced by BTZ was markedly diminished by SCH772984, indicated by the expression of proteasome subunit genes (Figure 6F). Together, our findings established that inhibition of ERK1/2, and in turn the NRF1-driven proteasome recovery response, can synergize with BTZ treatment to inhibit tumor growth *in vivo*.

**Figure 6.**
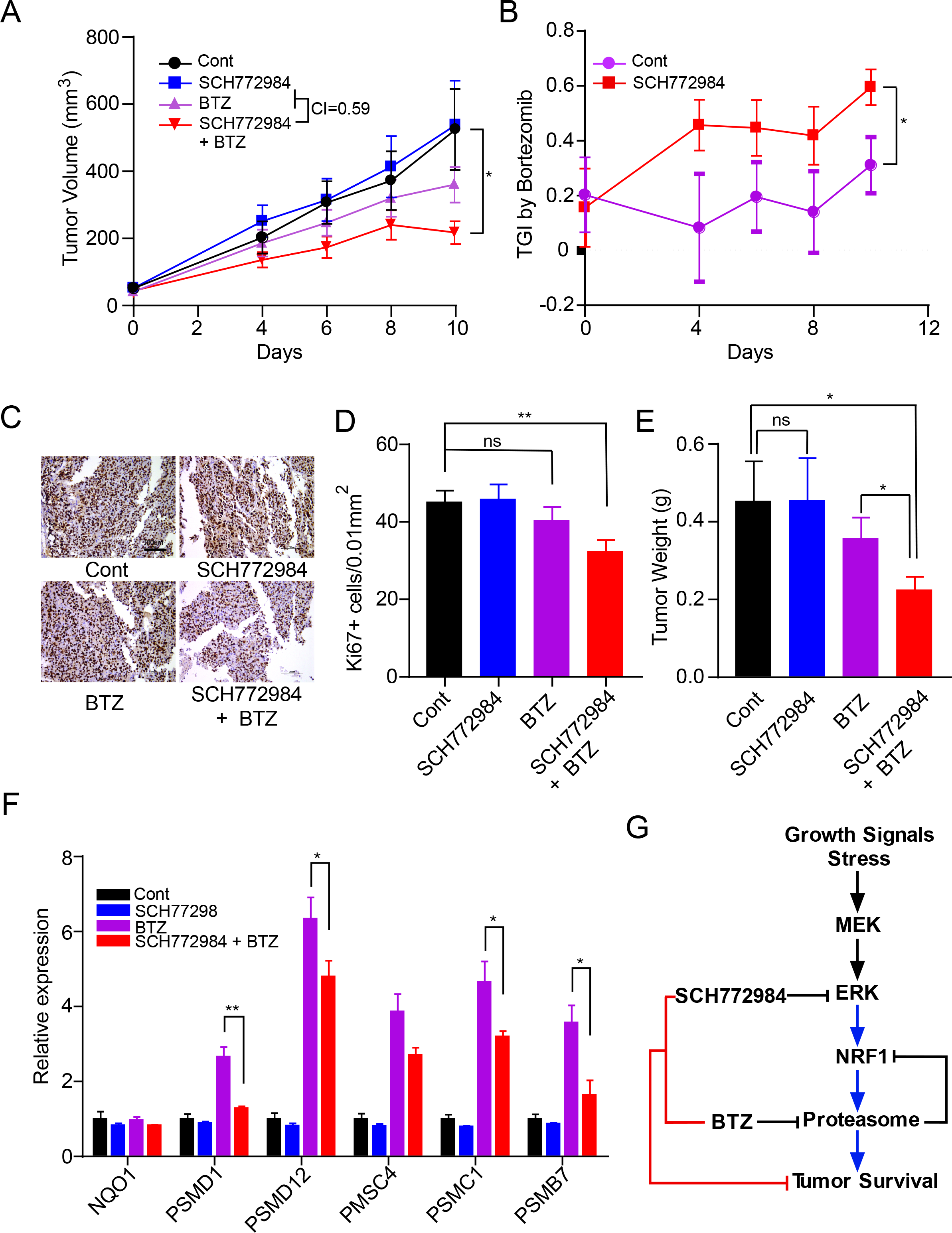
Proteasomal stress synergizes with ERK1/2 inhibition to suppress tumor growth *in vivo*. **A**. SCH772984 enhances BTZ-induced inhibition of A2058 tumor cell growth. Mice were treated with Control, SCH772984 (20 mg/kg, i.p. daily), BTZ (0.5 mg/kg, i.p. every other day), or a combination of SCH772984 and BTZ for 10 days, n=14 for each cohort. Tumor size was measured at day 0, 4, 6, 8 and 10, *p<0.05. **B**. Quantification of the tumor growth inhibitory (TGI) effect of BTZ alone (Cont) or BTZ combined with SCH772984, *p<0.05. **C, D**. Inhibition of tumor cell proliferation by combined BTZ and SCH772894 treatment. Representative images (C) and quantification (D) of Ki67 staining are shown for tumors collected from mice described in A (n=20 per cohort), Scale bar, 100 µm, ns, p>0.05, **p<0.01. **E**. Combined BTZ and SCH772894 treatment reduced tumor cell weight. A2058 tumors were weighed after a 10 day treatment with solvent control, SCH772984 (20 mg/kg, i.p. daily), BTZ (0.5 mg/kg, i.p. every other day), or a combination of SCH772984 and BTZ, ns, p>0.05, *p<0.05. **F**. Inhibition of the proteasome recovery response *in vivo* by ERK1/2 inhibition. An RT-PCR analysis shows the expression of 6 proteasome subunit genes in tumors collected in (A), *p<0.05, **p<0.01. **G**. Schematic depiction of the ERK1/2-NRF1 pathway and the proteasome recovery response. Inhibition of the ERK1/2-NRF1 pathway represents a promising strategy for pharmacologically enhancing the efficacy of proteasome inhibition in cancer therapy.

## Discussion

Through the evolutionarily conserved proteasome recovery response, NRF1/SKN-1A functions as the major transcriptional regulator of proteasome homeostasis (Deshaies, 2014; Lehrbach and Ruvkun, 2016; Li et al., 2011; Sha and Goldberg, 2014; Steffen et al., 2010). While it is clear that this response senses proteasome activity directly, it has remained largely unknown whether it might be induced or restricted by other regulatory inputs. Here we have determined that in *C. elegans* and humans, the proteasome recovery response depends upon signaling through the MAP kinase MPK-1/ERK1/2, which we have determined interacts with and phosphorylates NRF1, and is required for NRF1 nuclear localization. ERK inhibition is under investigation as a strategy of targeted combinatorial cancer therapy (Bryant et al., 2019; Kinsey et al., 2019), and we find that pharmacological targeting of this signaling dramatically enhances the tumor inhibitory effects of BTZ in an *in vivo* mouse model of human melanoma. Our results reveal an evolutionarily conserved coupling between the proteasome recovery response and mechanisms that stimulate ERK signaling, as well as a promising new strategy for enhancing the effectiveness of proteasome inhibition therapy in cancer.

ERK signaling can be stimulated by stress (Kyriakis and Avruch, 2012), and ERK activity appeared to be increased by BTZ treatment (Figure S5E). Phosphorylation of NRF1 by ERK was not required to maintain NRF1 levels or processing, but was increased by BTZ treatment (Figure 4B). This suggests that with respect to the proteasome recovery response ERK signaling may play a licensing role, rendering NRF1 nuclear localization and activity dependent upon signaling inputs that activate ERK1/2. Multiple peptide growth factors act through tyrosine kinase receptors to stimulate ERK signaling, which in turn regulates transcription factors involved in cell proliferation or differentiation (Roskoski Jr, 2012). We speculate that ERK phosphorylation of NRF1 might similarly link proteasome gene expression to growth signals, so that if these signals are inactive increased proteasome production is inhibited even if proteasome activity is low. Under low-growth conditions, this mechanism would limit accumulation of proteasome biomass, as well as potential proteasome-mediated increases in the free amino acid pool (Rousseau and Bertolotti, 2016; Zhang et al., 2014). Interestingly, in *S. cerevisiae* the ortholog of ERK1/2 and MPK-1 (Mpk1) enhances proteasome abundance by post-transcriptionally increasing expression of proteasome regulatory particle-assembly chaperones (RACs), and proteasome subunits (Rousseau and Bertolotti, 2016). Knockdown of the ERK family kinase ERK5 reduces RAC levels in human cells, suggesting that this pathway is conserved (Rousseau and Bertolotti, 2016). Our finding that MPK-1/ERK1/2 signaling is needed to license the metazoan SKN-1A/NRF1 proteasome recovery response reveals that the relationship between ERK/growth signaling and proteasome function is not only ancient but even broader, encompassing *C. elegans* and humans and extending to transcriptional regulation of the proteasome. Additionally, both NRF1 and its ortholog SKN-1A control many types of genes besides proteasome subunit genes (Hirotsu et al., 2012; Widenmaier et al., 2017) (Figure 2D and Table S2), and NRF1 has been implicated in metabolic functions (Li et al., 2011; Tsujita et al., 2014; Xu et al., 2005), suggesting that these functions are likely to be coupled to ERK and growth signaling as well.

Many aspects of NRF2 regulation and functions are understood in great depth (Cuadrado et al., 2019; Hayes and Dinkova-Kostova, 2014; Yamamoto et al., 2018), but work on NRF1 is at a comparatively early stage. It is an intriguing question how and why these two related proteins evolved in such a way that one of them (NRF1) is regulated post-transcriptionally through an elaborate process that involves ER-associated synthesis, processing, and degradation (Deshaies, 2014; Lehrbach et al., 2019). In *C. elegans* SKN-1A and SKN-1C correspond functionally to NRF1 and NRF2, respectively (Blackwell et al., 2015; Lehrbach and Ruvkun, 2016; Li et al., 2011; Oliveira et al., 2009) but are encoded by the same gene, and differ only by the presence of 90 N-terminal amino acids that mediate ER localization of the former (Figure S1A) (Lehrbach and Ruvkun, 2019). This suggests that NRF1/NRF2 and SKN-1 evolved from a common precursor that fulfilled many of their major functions, and that NRF1 and NRF2 likely arose from alternatively spliced isoforms of the same protein, an organization analogous to SKN-1 (Figure S1A). Remarkably, while the nuclear localization and activity of SKN-1C/NRF2 depends upon p38/PMK-1,2,3 signaling (Hourihan et al., 2016; Inoue et al., 2005), we find that SKN-1A/NRF1 function specifically requires ERK/MPK-1 signaling (Figure 6G). This suggests that the differential regulation of these two stress response factors by these respective MAPK pathways predated divergence of NRF1 and NRF2 into two distinct proteins. We speculate that the ER passage and processing events to which SKN-1A and NRF1 are subjected were critical for allowing SKN-1A/NRF1 and SKN-1C/NRF2 to be responsive to different stimuli. Furthermore, this control of SKN-1A/NRF1 at the ER and through the ERAD mechanism may allow it to be exquisitely sensitive to perturbations in proteasome activity.

To date, proteasome inhibition therapy has not been effective for treatment of solid tumors, or almost any other cancer aside from multiple myeloma, in which acquisition of resistance to proteasome inhibitors remains an intractable problem (Gandolfi et al., 2017; Manasanch and Orlowski, 2017). Nevertheless, it is clear that proteostasis could represent a point of sensitivity for many tumor types, for example including triple-negative breast cancer (Petrocca et al., 2013; Walerych et al., 2016; Weyburne et al., 2017). Our analysis of melanoma cells indicates that targeting of the NRF1 proteasomal recovery response through ERK1/2 inhibition represents a potentially valuable pharmacological option for augmenting the effectiveness of proteasomal inhibition therapy *in vivo*. ERK1/2 inhibition sensitizes additional cancer cell types to BTZ treatment *in vitro* (Figure 5C, D), suggesting that the results obtained *in vivo* with melanoma cells are likely to be applicable to other tumors. A2058 melanoma cells carry the V600E mutation in B-RAF, gain-of-function mutations in which are common genetic features in melanoma (Poulikakos et al., 2011). Our results indicate that interference with RAS-ERK signaling not only inhibits cell growth, but also confers the added benefit of inhibiting NRF1 and its response to proteasomal stress, suggesting that anti-proteasomal therapy should considered as a strategy for enhancing the effectiveness of inhibiting the ERK pathway. ERK is activated directly by MEK, which also phosphorylates and activates heat-shock factor (HSF1), and MEK inhibition has been shown to impair proteostasis and cooperate with BTZ to inhibit metastasis of A2058 cells in mice (Tang et al., 2015). Our results in *C. elegans* and human cells demonstrate that MEK signaling promotes proteostasis and resistance to BTZ not only through HSF1, but also through NRF1/SKN-1A (Figure 6G). Inhibition of MEK or ERK therefore provides two alternative adjuvant approaches to proteasome inhibition therapy, with one or the other potentially being more advantageous depending upon the tumor characteristics or toxicities encountered. Our findings also represent a proof of principle that pharmacological targeting of other nodes in NRF1 processing and the proteasome recovery response, including p97, DDI2, and NGLY1 (Figure 1A) could be promising therapeutic strategies (Figure 6G). Interference with components of this pathway from RAS to NRF1 might thus provide many options for developing tailored approaches to enhancing the effectiveness of proteasome inhibition therapy.

## Supplemental Figure Legends

**Figure S1.**
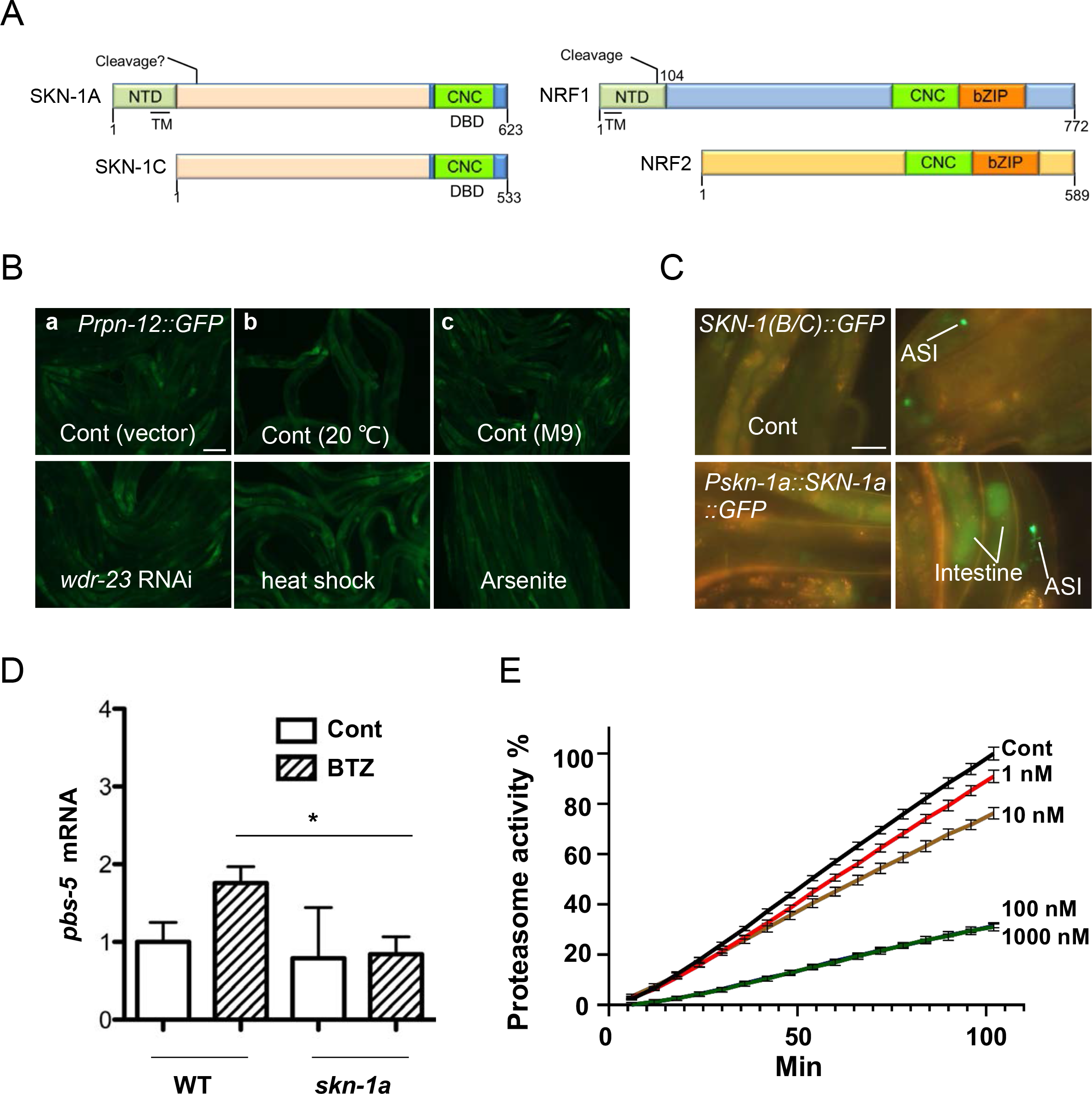
SKN-1A is required to regulate the proteasome recovery response in *C. elegans*. **A**. A schematic diagram comparing SKN-1A, SKN-1C, NRF1, and NRF2. The *skn-1a* mutation used in our experiments (*mg570*) (Lehrbach and Ruvkun, 2016) is located within the NTD and specifically eliminates expression of the SKN-1A isoform of SKN-1 (Figure S1A). NTD: N-terminal domain; TM: Transmembrane domain; CNC: Cap’n’Collar; bZIP: Basic leucine zipper; DBD: DNA-binding domain. **B**. The proteasome marker *Prpn-12::GFP* specifically responds to proteasomal stress. This reporter was not activated by multiple other stimuli, including knocking down *wdr-23*, (the inhibitor of SKN-1C) (Blackwell et al., 2015), heat shock, or treatment with the protein synthesis inhibitor cycloheximide or the ER stress inducer tunicamycin, Scale bar, 100 µm. **C**. BTZ increases expression of *Pskn-1a::SKN-1A::GFP* but not expression or nuclear accumulation of *SKN-1(B/C)::GFP*. Both *skn-1a* and *skn-1(b/c)* are constitutively expressed in the ASI neurons, while only SKN-1A is upregulated in the intestine by BTZ, Scale bar, 50 µm. **D**. BTZ-induced activation of the proteasome subunit gene *pbs-5* is dependent upon *skn-1a*, assayed by RT-PCR, *p<0.05. **E**. Inhibition of *C. elegans* proteasome activity by BTZ *in vitro*. Treatment with 100 nM BTZ was sufficient for significant inhibition, in contrast to the much higher levels required *in vivo* (see text).

**Figure S2.**
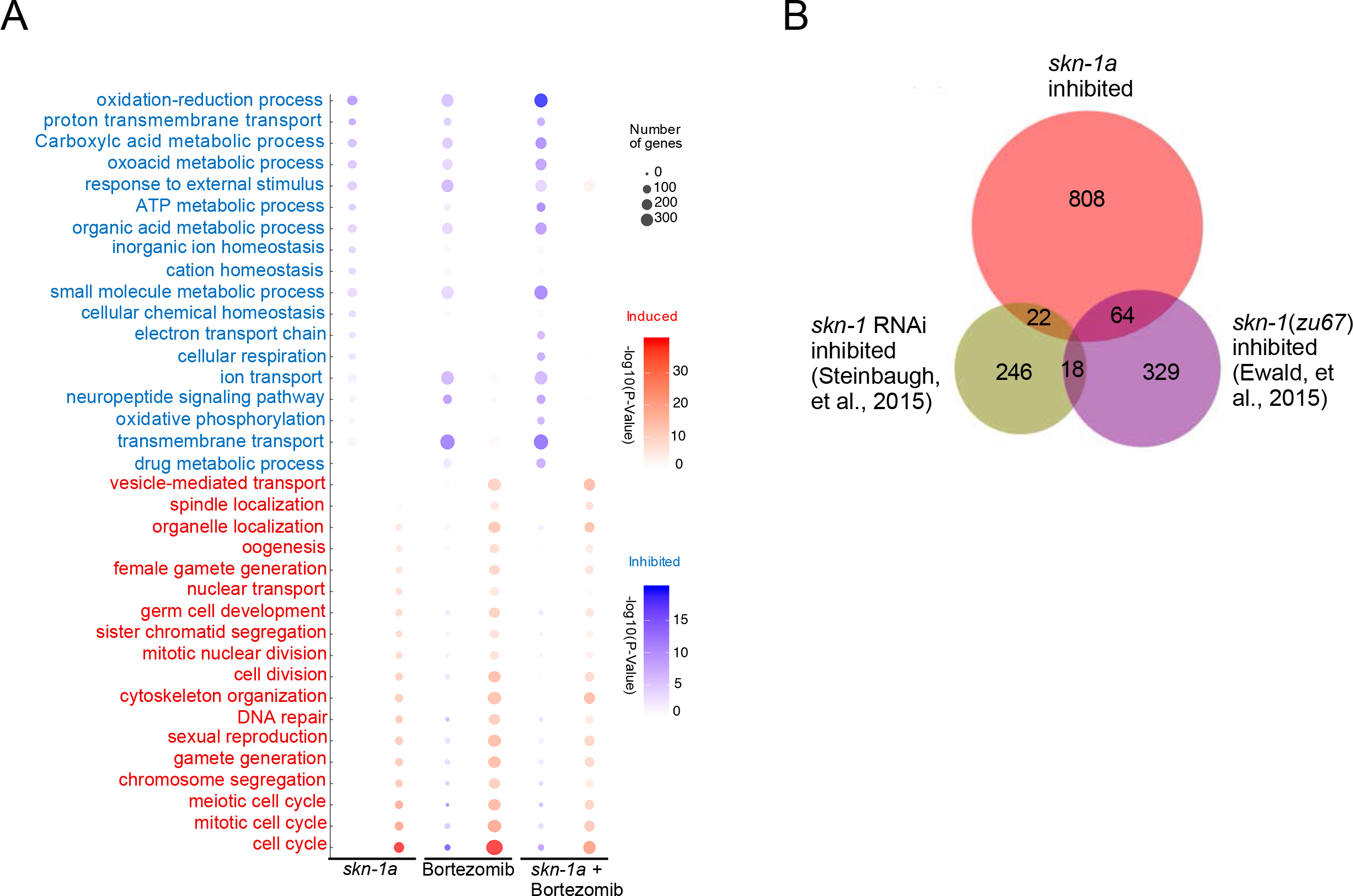
Regulation of proteasome subunit and multiple other gene categories by SKN-1A. **A**. GO term analysis of BTZ-induced and *skn-1a*-dependent genes. **B**. Overlap among genes that were downregulated in *skn-1a* mutants (this study), by *skn-1*(RNAi) in animals lacking germ cells (Steinbaugh et al., 2015) and by the *skn-1(zu67)* mutation in animals with reduced insulin-IGF-1 signaling (Ewald et al., 2015).

**Figure S3.**
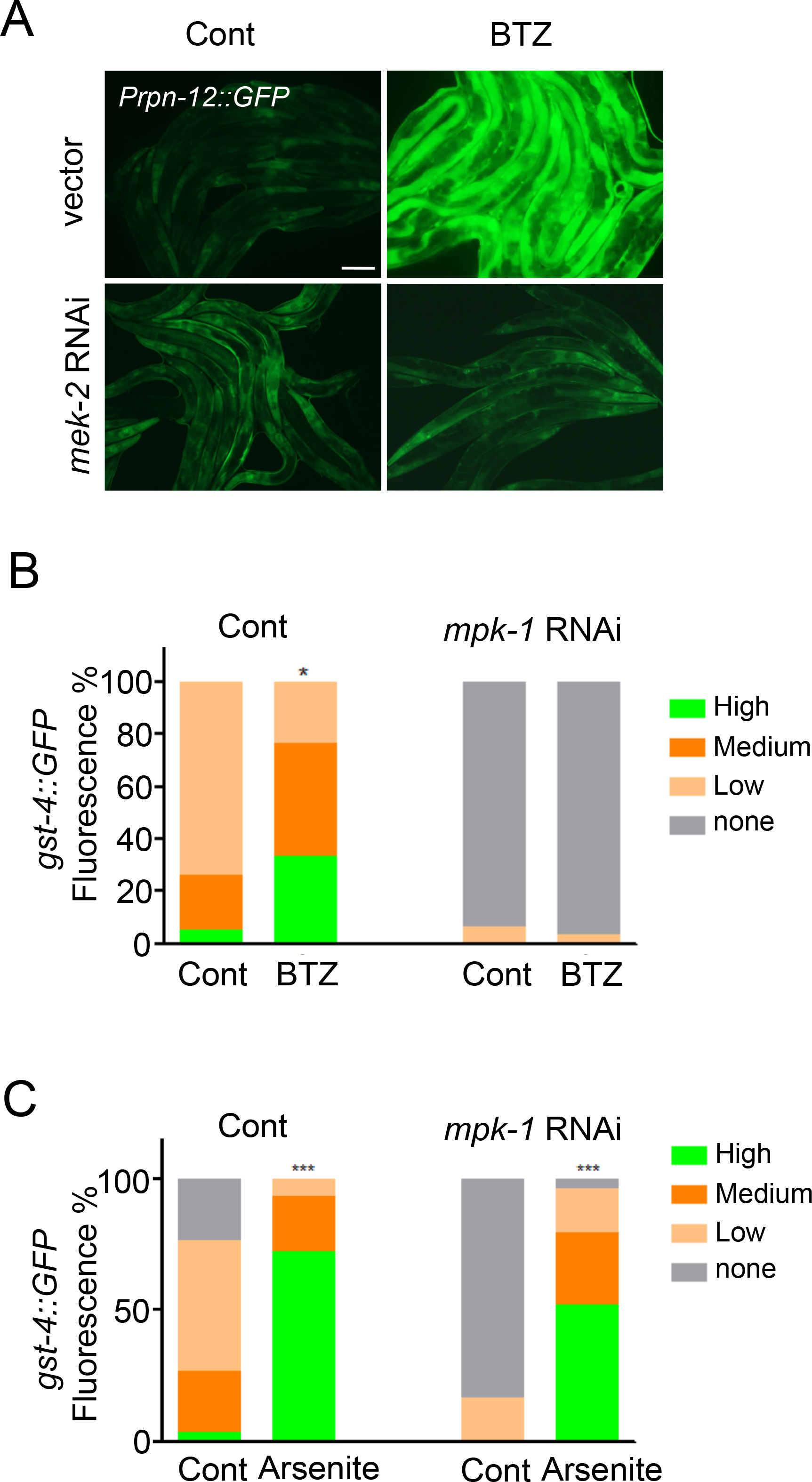
Dependence of the *C. elegans* proteasome recovery response on MPK-1/ERK signaling. **A**. *mek-2* RNAi blocks BTZ-induced expression of the *Prpn-12::GFP* reporter, and indicator of the proteasome recovery response. **B, C**. Quantification of *Pgst-4::GFP* activation by BTZ and Arsenite (oxidative) stress (refer to Figure 3D). Note specific inhibition of BTZ induction by *mpk-1* knockdown, *p<0.05, ***p<0.001.

**Figure S4.**
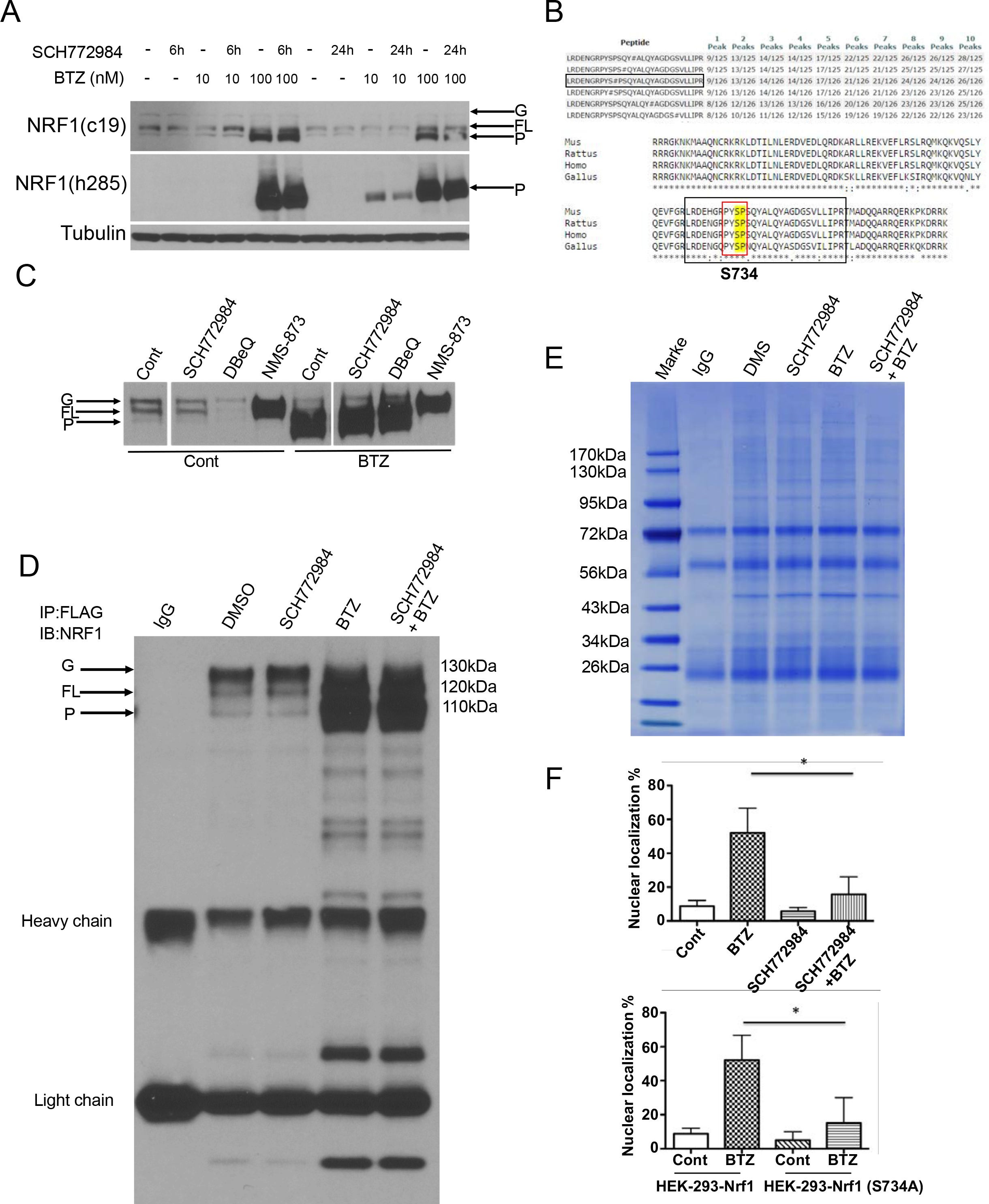
Human NRF1 nuclear localization depends upon phosphorylation by ERK1/2. **A**. Analysis of BTZ effects on NRF1 by two different antibodies. The c-19 antibody was used for further experiments in this study because it readily revealed all three forms of NRF1 (glycosylated, full length, and processed). **B**. A phosphorylated peptide that included the predicted ERK1/2 motif PYSP was identified by mass spectrometry at the C terminal of NRF1 (see text). **C**. In HEK293-NRF1^3×FLAG^ cells, inhibition of P97/VCP ATPase by the P97 inhibitors DBeQ and NMS-873 results in accumulation of the glycosylated (Radhakrishnan et al., 2014) but not further processed forms of NRF1. By contrast, ERK1/2 inhibition with SCH772984 does not disrupt NRF1 processing. **D**. Western blot detection of FLAG-tagged NRF1 by NRF1 antibody in HEK293-NRF1^3×FLAG^ cells. **E**. Colloidal blue staining of the co-immunoprecipitation sample that was analyzed by mass spectrometry. **F**. Quantification of NRF1 nuclear localization in the experiments shown in Figures 4D and 4E, *p<0.05.

**Figure S5.**
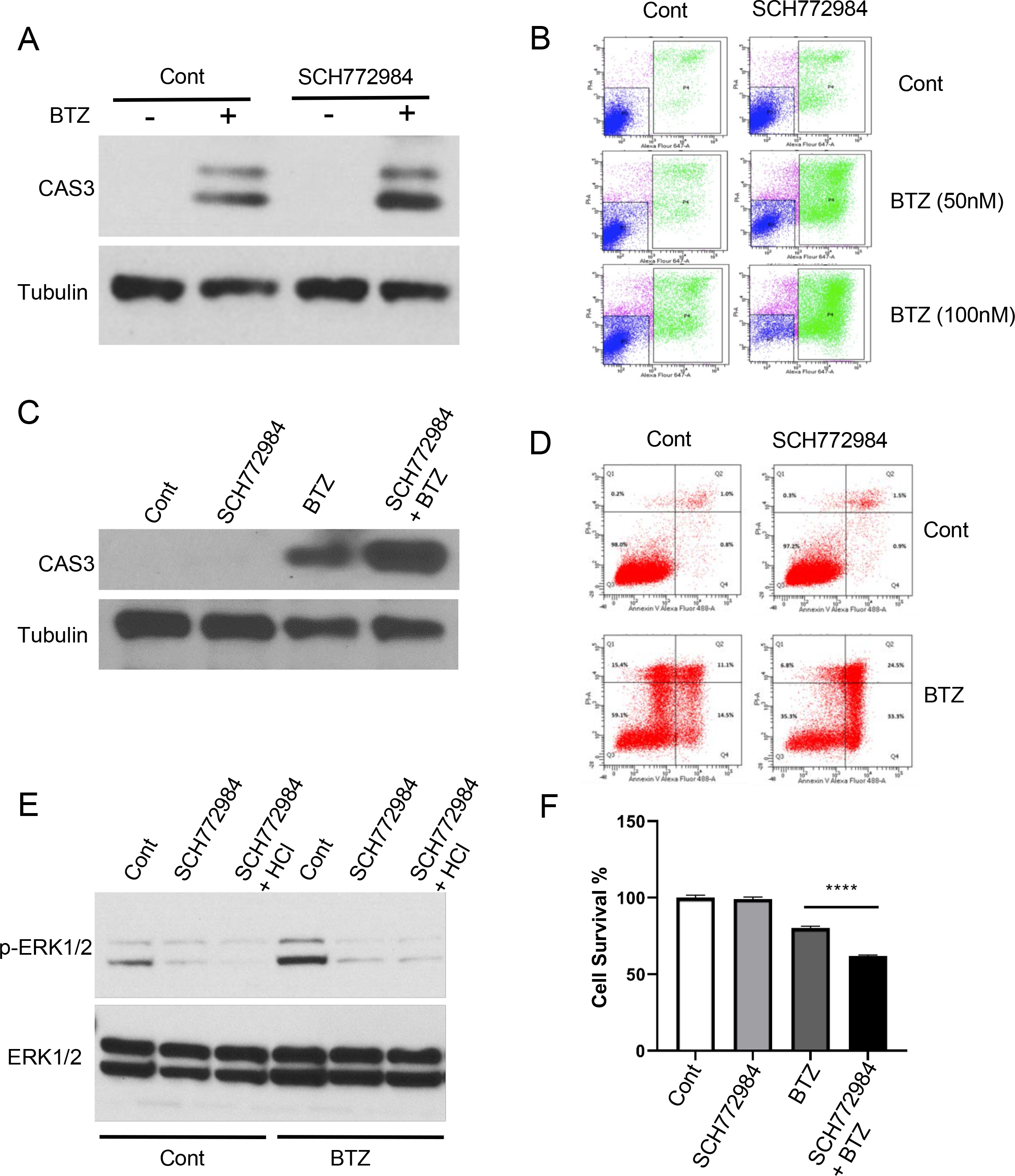
ERK1/2 inhibition sensitizes human tumor cells to proteasomal stress. **A**. SCH772984 treatment increases BTZ-induced cell death in the Melanoma cell line A2058, detected by CAS3 cleavage. **B**. Flow cytometry detection of A2058 cell apoptosis (monitored by propidium iodide and Annexin V staining) after treatment with BTZ or/and SCH772984. Quantification of the Annexin V (+) PI (-) fractions are shown in Figure 5E. **C**. SCH772984 treatment increases BTZ-induced apoptosis in MM.1s cells, detected by western blotting for CAS3 cleavage. **D**. Flow cytometry analysis of apoptosis (assayed by propidium iodide and Annexin V staining) in MM.1s cells that were treated with BTZ or/and SCH772984. Quantification of the Annexin V (+) PI (-) fraction is shown in Figure 5D. **E**. ERK1/2 phosphorylation, an indicator of activity, is diminished by treatment with the ERK1/2 inhibitors SCH772984 and SCH772984-HCL. **F**. MM.1s cell growth and viability is impaired by combined BTZ and SCH772984 treatment, indicated by the Cell Counting Kit-8 colorimetric assay of cell proliferation, ****p<0.0001.

**Figure S6.**
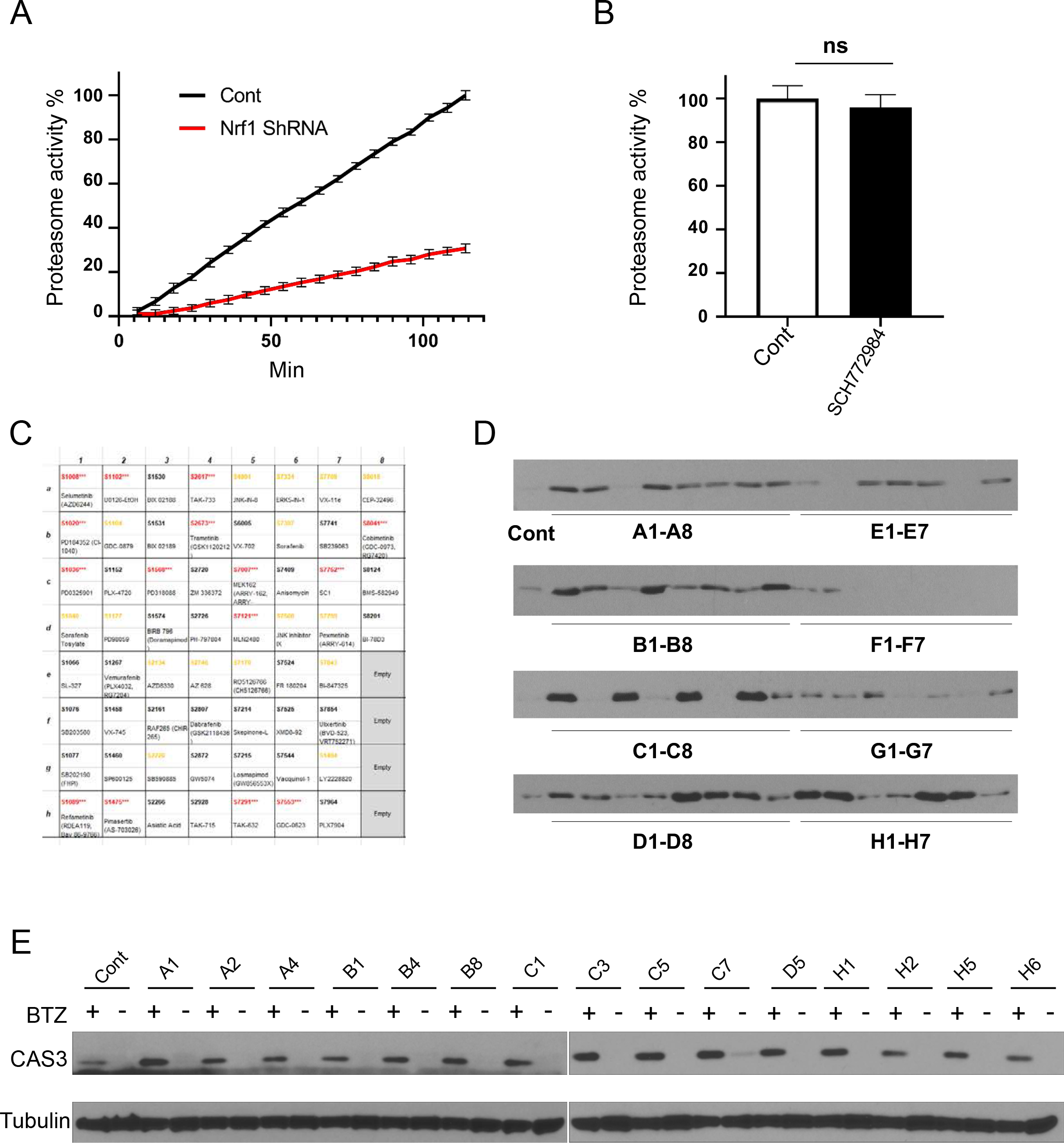
Small molecule screen to identify compounds that reduce BTZ resistance. **A**. Proteasome activity was reduced in SH-SY5Y Nrf1 shRNA cells compared to SH-SY5Y shRNA control cells, ****p<0.0001. **B**. SCH772984 treatment does not significantly reduce the background level of proteasome activity in HepG2 cells, ns, p>0.05. **C**. Screening of a commercial MAPK inhibitor library for small molecules that could increase the level of BTZ-induced apoptosis in HepG2 cells, as shown in D. Red and yellow indicate strong (robust CAS3 increase over control) and moderate (detectable CAS3 increase over control) hits, respectively. **D**. Induction of apoptosis in HepG2 cells that were treated with BTZ along with the indicated MAPK inhibitors from the library in C, with cell death detected by western blotting for cleavage of CAS3. **E**. Repeat analysis of “strong” hits from the screen shown in C and D.

**Figure S7.**
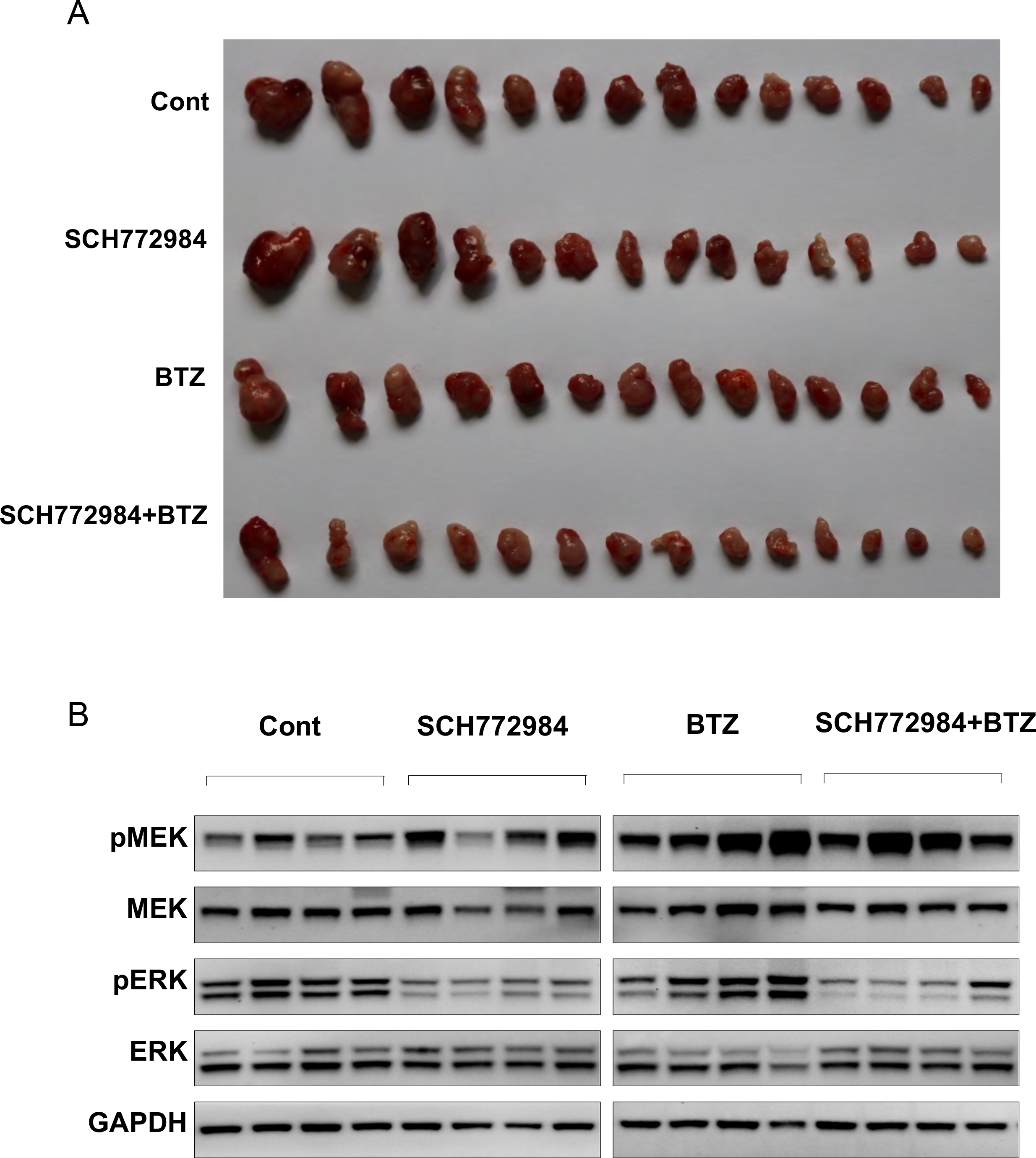
ERK1/2 inhibition synergizes with proteasomal stress to suppress melanoma tumor growth *in vivo*. **A**. Inhibition of A2058 tumor growth by SCH772984 and BTZ treatment *in vivo*. A2058 cells were injected into nude mice, and treated with solvent control, SCH772984 210 mg/kg, i.p. daily), BTZ (0.5 mg/kg, i.p. every other day), or a combination of SCH772984 and BTZ for 10 days. All tumors of each cohort were collected and shown. **B**. Western blot showing the relative levels of MEK, pMEK, ERK and pERK in tumor samples from (A). SCH772984 inhibits ERK but not MEK activity.

## STAR★METHODS

Detailed methods are provided in the online version of this paper and include the following:

- KEY RESOURCES TABLE
- LEAD CONTACT AND MATERIALS AVAILABILITY
- EXPERIMENTAL MODEL AND SUBJECT DETAILS
  ➢ *C. elegans* strains, cell lines and reagents
- METHOD DETAILS
  ➢ High throughout ORF screening
  ➢ BTZ tolerance assays
  ➢ Quantitative RT–PCR
  ➢ RNA sequencing experiments
  ➢ Immunofluorescence
  ➢ Immunoblot and immunoprecipitation
  ➢ Phosphorylation mass spectrometry
  ➢ Cellular viability assays
  ➢ Flow cytometry
  ➢ Proteasome activity assay
  ➢ Small molecules screening
- DATA AND SOFTWARE AVAILABILITY
- ADDITIONAL RESOURCES

## SUPPLEMENTAL INFORMATION

Supplemental Information can be found online at ……

## ACKNOWLEDGEMENTS

We thank Raymond Deshaies, Fred Goldberg, Ken Anderson, and Gary Ruvkun for reagents, and Fred Goldberg, Gary Ruvkun, Nic Lehrbach, Ken Anderson, and Giada Bianchi for helpful discussions. The work was funded by support to TKB from the NIH (R35 GM122610 and R01 AG054215) and an NIDDK DRC grant to the Joslin Diabetes Center (P30 DK036836). PZ was supported by an NIH T32 training award (T32 DK007260) and a fellowship from the Iacocca Family Foundation. YXF was supported by NFSC grants (31871369, D19H160009, and 2019R01001). HN and AJMW are supported by NIH grant R35GM122502. Some strains were provided by the CGC, which is funded by the NIH (P40 OD010440).

## AUTHOR CONTRIBUTIONS

Conceptualization, PZ and TKB; Methodology, PZ; Investigation, PZ, HYQ, HN, JH, FZ, ZW, FZ, MI; Writing – Original Draft, PZ and TKB; Writing – Review & Editing, PZ, YXF, AJMW, MI, and TKB; Supervision, TKB, YXF and AJMW; Funding Acquisition, PZ, AW, AJMW, YXF and TKB.

## STAR★METHODS

### KEY RESOURCES TABLE

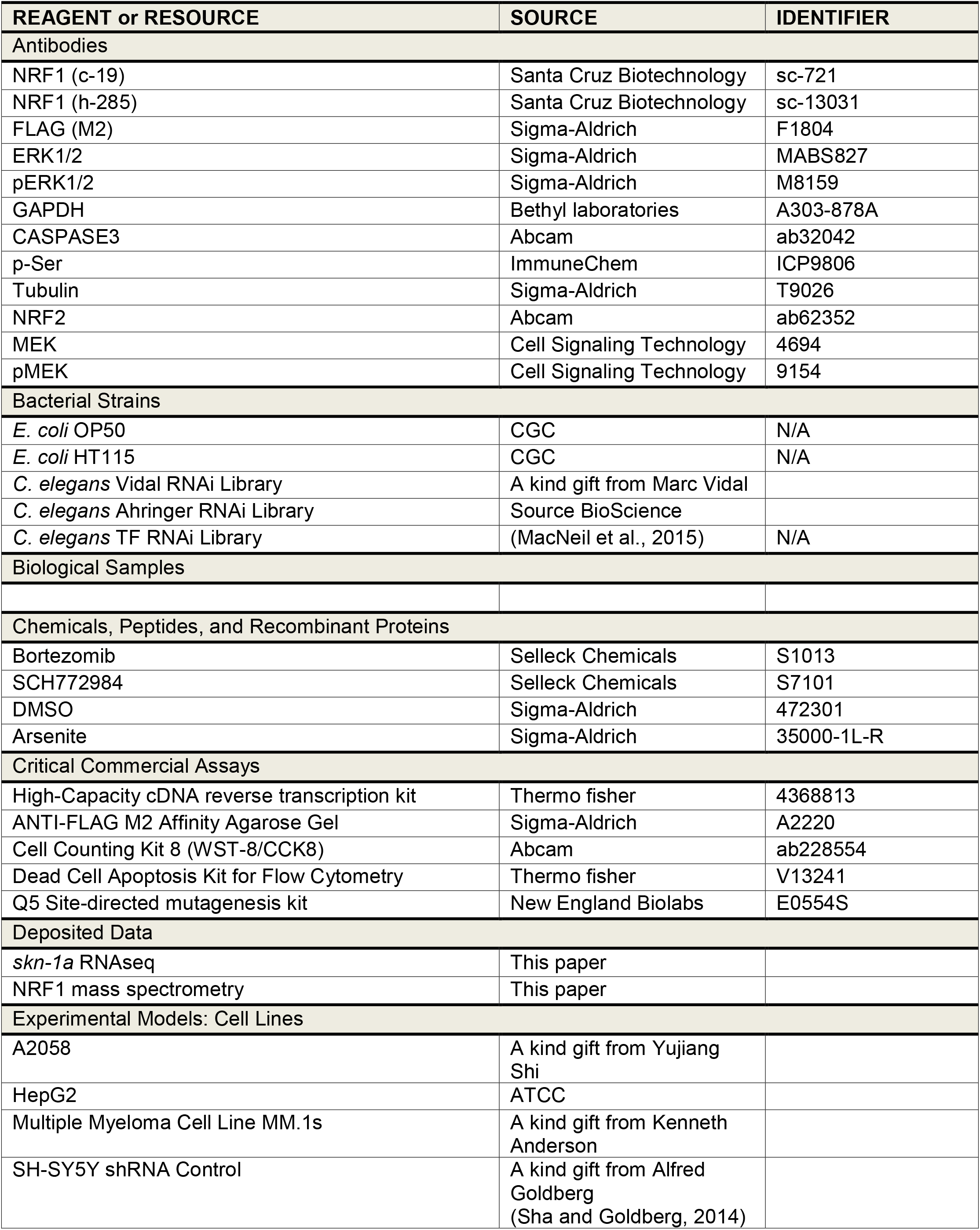
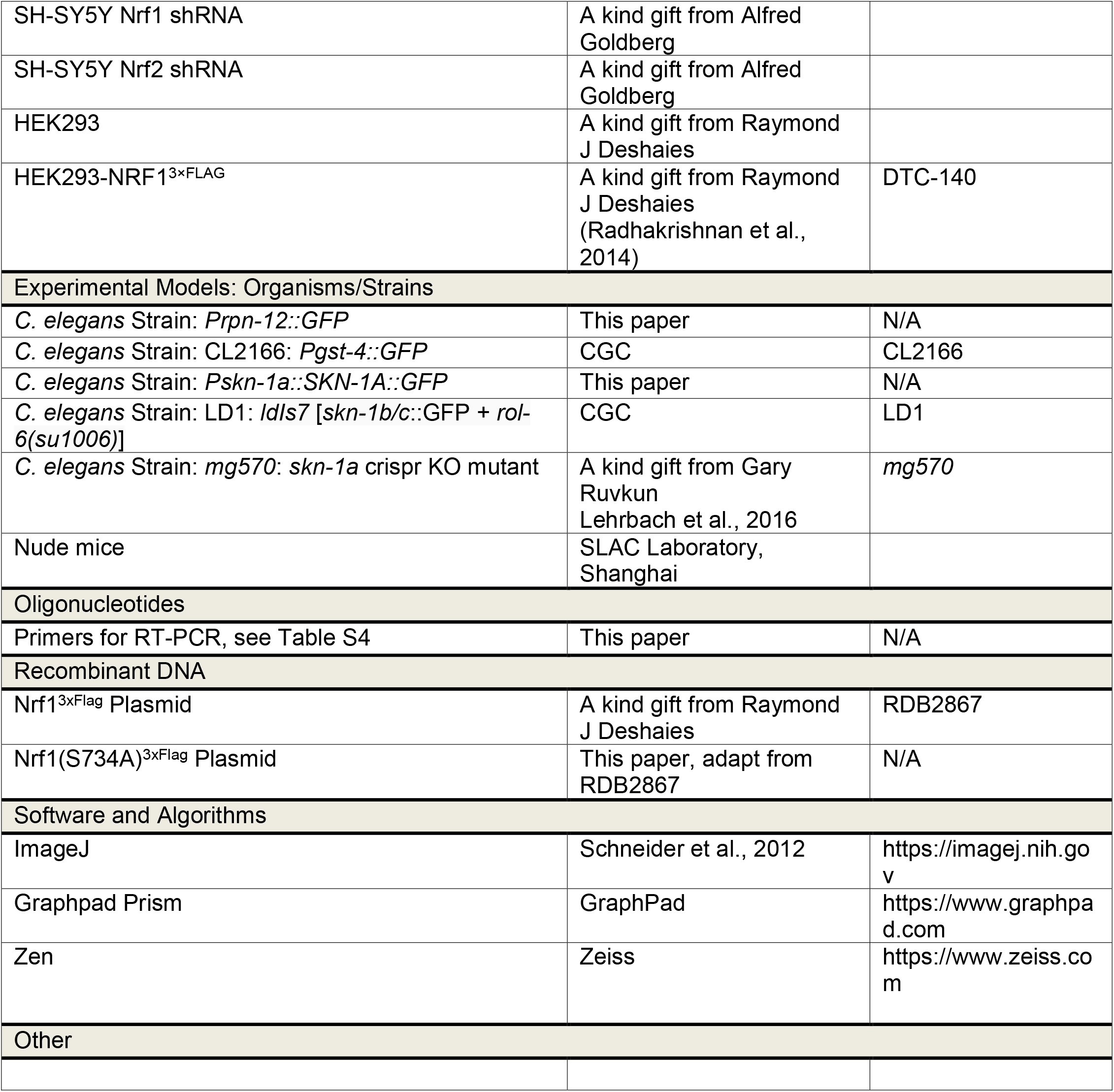

### CONTACT FOR REAGENT AND RESOURCE SHARING

Further information and requests for resources should be directed to and will be fulfilled by the Lead Contact, T Keith Blackwell.

## EXPERIMENTAL MODEL AND SUBJECT DETAILS

### *C. elegans* strains, cell lines and reagents

*C. elegans* were maintained on standard media at 20°C and fed *E. coli* OP50. RNAi was performed as described in (Kamath and Ahringer, 2003). The *Prpn-12::GFP* integrated transgene was generated and outcrossed 4 times in our lab. Some strains were provided by the CGC. *skn-1a*(*mg570)* (Lehrbach and Ruvkun, 2016) was kindly provided by Gary Ruvkun.

MM.1s cells and other multiple myeloma cell lines were kindly provided by Kenneth Anderson, HEK293-NRF1^3xFLAG^ cell lines by Raymond J Deshaies, and SH-SY5Y Nrf1 ShRNA and Nrf2 ShRNA cell lines by Alfred Goldberg. Multiple Myeloma cell lines were grown in RPMI (Cellgro), 10% FBS and 1% penicillin/streptomycin. HepG2, HEK293 and SH-SY5Y cells were grown in DMEM (Cellgro), 10% FBS and 1% penicillin/streptomycin.

BTZ, SCH772984, and the MAPK inhibitor library were purchased from Selleck Chemicals. All Chemicals were diluted with DMSO. The concentration of BTZ for treatment of cell lines was 100 nM, except for Multiple myeloma cell lines (10 nM). The concentration of MAPK inhibitors and SCH772984 used for cell line treatments was 100 nM. Cells were treated with BTZ for 18 hours with or without 6 hour a pretreatment of MAPK inhibitors or SCH772984.

## METHOD DETAILS

### High throughout ORF screening

The *Prpn-12::GFP* strain was plated in 96-well plates containing RNAi bacteria at ∼20 L1 worms per well. After 48 hours, fluorescence was examined on a Zeiss M2 microscope. At that time 100 µL M9 containing 50 µM BTZ (two replicates) was added to each well, then after 24 hours fluorescence was reexamined on the Zeiss M2 microscope. The entire screening experiment was performed twice.

### BTZ tolerance assays

L4 stage worms were seeded into 96-well plates (20 worms per well). Twenty-four hours after seeding, 500 µM BTZ was added, with the final volume of DMSO not exceeding 1%. Worms were incubated for 24-72 h following addition of BTZ. Worm viability was measured and calculated as a percentage of control (untreated worms) after background subtraction. A minimum of six replicates were performed for each worm and drug combination.

### Quantitative RT–PCR

mRNA was extracted from worms (∼100) and cells (10^5^∼10^6^) using the RNeasy kit (Qiagen). 1 µg mRNA was used for subsequent reverse transcription using the SuperScript III First-Strand Synthesis SuperMix (Invitrogen). The reverse transcription reaction (5 µl) was used for quantitative PCR using SYBR Green PCR Master Mix and gene-specific primers, in triplicate, using an ABI 7700 Real Time PCR System. Oligonucleotide primers used are listed in Table S4.

### RNA sequencing experiments

For each RNAseq condition, three independent biological replicates that had been obtained in parallel were examined. For each sample, 2,000 synchronized N2 or *skn-1a* L1 worms were grown at 20°C in a 10 cm plate. After 48 hours, when they reached the L4 stage, each group was placed in 10 mL M9 buffer or 50 µM BTZ in M9 buffer for 24 hours. They were then collected with M9 buffer into a 1.5 mL Eppendorf tube, and washed 4 times in M9. 300 µL TRIzol (Sigma-Aldrich, St. Louis, MO) was added to each tube, and the tubes were frozen in −80°C. Total RNA was extracted using TRIzol, and purified with Direct-zol RNA Kits (Zymo Research, Irvine, CA). Purified RNA samples were DNase treated before sending for sequencing. The quantity and quality of isolated RNA was quantified with a Qubit (1.0) Fluorometer (Life Technologies) and a Bioanalyzer 2100 (Agilent, Santa Clara, CA, USA). The preparation of RNAseq libraries were performed using the TruSeq Stranded mRNA Sample Prep Kit (Illumina, San Diego, CA, USA) at Novogene Co., Ltd. Paired-end reads (150 bp) were generated on the Illumina platform. FASTQ output files were aligned to *C. elegans* WBcel269 reference genome using STAR with a 2-pass procedure (Dobin et al., 2013). The annotated genes and transcripts were quantified using RSEM (Li and Dewey, 2011). Differential expression analysis of genes and transcripts were performed in R using the Bioconductor package DESeq2 (Love et al., 2014). A threshold of FDR (padj) < 0.01 was set to determine differentially expressed targets.

### Immunofluorescence

NRF1^3xFLAG^ was overexpressed in HEK293 cells (Radhakrishnan, den Besten et al. 2014), and its expression was detected with mouse anti-FLAG antibody and the corresponding anti-IgG antibody conjugated to Alexa 594. Cells were mounted onto glass slides using Immu-Mount (Shandon) containing 4,6-diamidino-2-phenylindole (DAPI) to stain nuclei. Confocal laser microscopy was performed on a Zeiss LSM 510 microscope. Quantitative analysis of immunofluorescence data was carried out by histogram analysis of the fluorescence nuclear localization of 10 random images by 2 independent replicates. The results of the analysis of 10 images acquired in each experimental condition were then combined to allow quantitative estimates of changes in NRF1 localization.

### Immunoblot and immunoprecipitation

Cells were washed twice with ice-cold PBS and lysed with 1% NP-40 buffer (150 mM NaCl, 50 mM Tris pH 7.5, 2 mM EDTA pH 8, 25 mM NaF and 1% NP-40) containing protease inhibitors (Roche). Lysates were quantified (Bradford assay), normalized, reduced, denatured (95 °C) and resolved by SDS gel electrophoresis on 10% Tris/Glycine gels. Protein was transferred to PVDF membranes and probed with primary antibodies recognizing pERK1/2 (T202/Y204), CAS3 (Cell Signaling Technology; 1:1,000), Actin (1:1,000). After incubation with the appropriate secondary antibody (anti-rabbit, anti-mouse IgG, 1:10,000 dilution), proteins were detected using chemiluminescence (Pierce).

Immunoprecipitations were performed overnight at 4 °C in 1% NP-40 lysis buffer, as described above, at a concentration of 1 µg/µl total protein using an ANTI-FLAG M2 Affinity Agarose Gel (Sigma). Beads were centrifuged and washed three times in lysis buffer and eluted and denatured (95 °C) in 2X reduced sample buffer (Invitrogen). Immunoblots were performed as above.

### Phosphorylation mass spectrometry

Three biological replicate experiments were performed in this analysis. After 18 h BTZ treatment, cells were collected and sonicated. After centrifugation, lysates were quantified (Bradford assay), normalized, reduced, denatured (95 °C) and resolved by SDS gel electrophoresis on 10% Tris/Glycine gels, and staining using a Colloidal Blue Staining Kit (Invitrogen). Protein bands in the range of 90-130 kDa were excised and sent for Phosphorylation mass spectrometry analyses at the Taplin Mass Spectrometry Facility of Harvard University, which was performed as described in (Wu, Haas et al. 2011).

### Cellular viability assays

MM.1s cells were treated with BTZ and/or SCH772984 as described above. Following selection, cells were plated (1 × 10^5^ cells per well) onto a 96-well plate in triplicate. Viable cells were counted using the wst-8 Kit and a Microplate Reader, following manufacturer’s specification. Triplicate cell counts were averaged and normalized relative to that of the control cell.

### Flow cytometry

Approximately 1×10^6^ MM.1s or A2058 cells were seeded on a 6-well plate overnight, then were treated with 10 nM BTZ with or without 6 hour pretreatment of 100 nM SCH772984. The cells were next spun down at 1000 g for 5 min, and washed with PBS for three times. The cell pellets were resuspended in 250 µl binding buffer (0.01 M HEPES, pH 7.4, containing 140 mM NaCl and 25 mM CaCl2) containing 2.5 µl Annexin V-FITC (BD Phamingen, CA, USA) and 0.5 µl PI (6 mg/ml) (Sigma). The cells were incubated at room temperature in the dark for 20 min. Then the cells were analyzed using a BD LSRFortessa cell analyzer (BD Biosciences). The signal was detected by BD LSRFortessa Blue Laser (FITC: ex 494 nm/em 519 nm, PE-Texas Red: ex 488 nm/em 615 nm, PerCP-Cy5.5: ex 482 nm/em 695 nm).

### Proteasome activity assay

Worms or cells were Sonicated in ice-cold buffer (10% glycerol, 25 mMTris/HCl pH 7.4, 10 mM MgCl_2_, 4 mM ATP and 1 mM dithiothreitol and centrifuged at 12000 g for 15 min at 4°C to remove debris. Protein concentrations were determined by the Bradford assay, using BSA as the protein standard. Chymotrypsin-like activity was determined by a fluorometric assay, using Suc–Leu–Leu–Val–Try–7-amino-4-methylcoumarin (Suc–LLVY–AMC) (Enzo Life Sciences) as the substrate (Sha and Goldberg 2014). Reactions were initiated by adding 50 μM substrate followed by incubation at 37°C for 30 min. Release of fluorogenic AMC was monitored at 360 nm excitation and 460 nm emission using a fluorometric plate reader.

### In vivo tumor growth assay

A2058 cells (1×10^6^ cells/site) were injected into both flanks of male nude mice (6-8 weeks age). Tumors were measured every two days with a caliper, and the tumor volume was calculated using the formula: volume = 0.5 × length × width^2^. At the endpoint, mice were euthanized and dissected. Xenografts were weighted after the dissection. The results are presented as the mean ± SE.

For the *in vivo* drug treatment assay, A2058 tumor-bearing mice were randomized into four cohorts once the size of tumor reached 50 mm^3^. The mice were treated with solvent control (PBS), SCH772984 (20 mg/kg, i.p. injection, once daily), Bortezomib (0.5 mg/kg, i.p. injection, once every two days), or a combination of SCH772984 and Bortezomib for 10 days. The tumor measurement and dissection were performed as mentioned above. Mice were cared for in accordance with guidelines approved by the First Affiliated Hospital, Zhejiang University School of Medicine.

### Small molecule screening

100 µL HepG2 cells (1 × 10^6^) were seeded in 96-well plates for 24 h. 10 µM MAPK inhibitors from the library (Selleckchem) were added at a 1:100 dilution. Six hours after pretreatment, DMSO (1:100) or 100× BTZ (10 µM in DMSO) was added to a final concentration of 100 nM. Cell viability was detected by CAS3 Western Blot, 18 hour after the addition of BTZ.

## ADDITIONAL RESOURCES

None.

